# Elevated HLA-E and NKG2A as a Consequence of Chronic Immune Activation Defines Resistance to *M. bovis* BCG Immunotherapy in Non-Muscle-Invasive Bladder Cancer

**DOI:** 10.1101/2022.03.06.483198

**Authors:** D. Ranti, Y.A. Wang, J. Daza, C. Bieber, B. Salomé, E. Merritt, J.A. Cavallo, E. Hegewisch-Solloa, E.M. Mace, A.M. Farkas, S. Shroff, F. Patel, M. Tran, T. Strandgaard, S. V. Lindskrog, L. Dyrskjøt, J. Qi, M. Patel, D. Geanon, G. Kelly, R.M. de Real, B. Lee, S. Kim-Schulze, T.H. Thin, M. Garcia-Barros, K.G. Beaumont, H. Ravichandran, Y. Hu, Y-C. Wang, L. Wang, D. LaRoche, Y. lee, R.P. Sebra, R. Brody, O. Elemento, A. Tocheva, B. D. Hopkins, P. Wiklund, J. Zhu, M.D. Galsky, N. Bhardwaj, J.P. Sfakianos, A. Horowitz

## Abstract

*Mycobacterium bovis* Bacillus Calmette-Guerin (BCG), the first-line treatment for non-muscle invasive bladder cancer (NMIBC), promotes the production of inflammatory cytokines, particularly interferon (IFN)-γ. Prolonged inflammation and IFN-γ exposure are known to cause an adaptive immune response, enabling immune escape and proliferation by tumor cells. We investigated HLA-E and NKG2A, a novel T and NK cell checkpoint pathway, as a driver of adaptive resistance in BCG unresponsive NMIBC. We observed ubiquitous inflammation in all patients after BCG immunotherapy, regardless of recurrence status. IFN-γ was shown to drive tumor expression of HLA-E and PD-L1. Further, NKG2A-expressing NK and CD8 T cells were enriched in BCG unresponsive tumors and with enhanced capacity for cytolytic functions. Strikingly, *in situ* spatial analyses revealed that HLA-E^HIGH^ tumors are activated to recruit NK and T cells via chemokine production, potentially sparing HLA-E^LOW^ tumors that would otherwise be susceptible to lysis. Finally, blood-derived NK cells retained anti-tumor functions at the time of tumor recurrence. These data support combined NKG2A and PD-L1 blockade for BCG unresponsive disease.

## Introduction

The only FDA-approved, first-line treatment for high-risk non-muscle-invasive bladder cancer (NMIBC) is intravesical *Mycobacterium bovis* Bacillus Calmette-Guerin (BCG), which is hypothesized to act in part via the release of inflammatory cytokines, notably IFN-γ (1). Tumor recurrence following BCG immunotherapy is common, with local recurrence rates between 32.6 to 42.1% and progression rates between 9.5 to 13.4% (2). Radical cystectomy is the only definitive treatment option for BCG-unresponsive disease, leaving patients at high risk for complications and diminished quality of life (3,4). Currently, there is substantial interest and numerous ongoing clinical trials to identify novel therapeutic options for BCG-unresponsive NMIBC.

Despite BCG’s use as NMIBC’s first-line immunotherapy for over 30 years, the mechanisms underlying tumor evasion of BCG’s therapeutic benefits are poorly understood. BCG’s six-dose induction regimen was devised anecdotally based on the packaging of BCG used in Morales et al.’s seminal study (5). Only a single dosing study has been performed to assess the immunologic response with this dosing regimen, which concluded that six doses stimulated an immune response but did not investigate the ramifications of the magnitude nor the duration of that immune response (6). Contributing to the difficulty in predicting BCG response is the variety of immune lineages implicated in its success, including neutrophils, monocytes, macrophages, dendritic cells (DCs), T cells, and natural killer (NK) cells (7).

Adaptive immune resistance, in which cancers are recognized by immune cells, but adaptive changes prevent effective clearance, is an important mechanism of therapeutic resistance (8). IFN-γ, a prominently BCG-induced cytokine, has been shown to have both pro- and anti-tumorigenic functions across an array of tumor indications (9). The equilibrium established by IFN-γ signaling may be critical in preventing therapeutic resistance. In melanoma and lung cancer, dysfunctional and overactive IFN-γ signaling is known to drive T cell exhaustion via upregulation of PD-L1 (10). In settings of chronic infection, a host-target stalemate can occur due to persistently elevated interferon signaling, the blockage of which can reduce viral burden (11–13). The impact of long-term intravesical BCG, which leads to immune stimulation via IFN-γ secretion, and its role in immune dysregulation and exhaustion in BCG unresponsive NMIBC is currently unknown.

Previous work comparing BCG-unresponsive to pre-treatment tumors showed a significant increase in PD-L1 expression on unresponsive samples (14). Immunotherapeutic strategies, especially those interrupting the PD-1/PD-L1 axis to unleash exhausted immune cells, have improved NMIBC treatment. Unfortunately, PD-1 response rates in NMIBC remain low, at ~19% durable response at 12 months (15–18). Further, PD-L1 expression levels alone have failed to accurately predict response to immunotherapy in NMIBC, suggesting that alternative mechanisms of immune suppression are important (19).

Recently, the HLA-E/NKG2A axis has been identified as a potent immune checkpoint regulating both CD8^+^ T and NK cells (20–22). Abrogation of NKG2A on NK cells increased effector activity against tumor cells and potentiated anti-tumor function (20,22) and blocking NKG2A on CD8 T cells in a murine model increased tumor vaccine response, even in a PD-1 resistant setting (20,22). Important to BCG immunotherapy, HLA-E has been shown to increase in the setting of IFN-γ release (23). In this study, we pose a novel hypothesis that BCG induces changes in the bladder tumor microenvironment (TME), including expression of NKG2A and PD-1 on CD8^+^ T cells, NKG2A on NK cells, and HLA-E and PD-L1 on tumor cells, and these phenotypic changes are mechanistically linked to treatment resistance and tumor recurrence. This study investigates HLA-E and NKG2A as a novel axis of resistance induced by chronic inflammation and suggests a potential combination strategy in NKG2A and PD-L1 blockade for overcoming resistance to BCG immunotherapy for treating NMIBC.

## Results

### Clinical cohort description

Cohort descriptions of patient samples used for the below experiments are described in Table 1. Patient samples were collected and processed at The Mount Sinai Hospital, and validation cohorts were collected and processed at Aarhus University. Median patient age ranged from 64 – 70.7 years old; median months to recurrence ranged from 1.7 to 2.9 months; median months to progression ranged from 4.43 to 23.28 (Table 1).

**Table 1:**
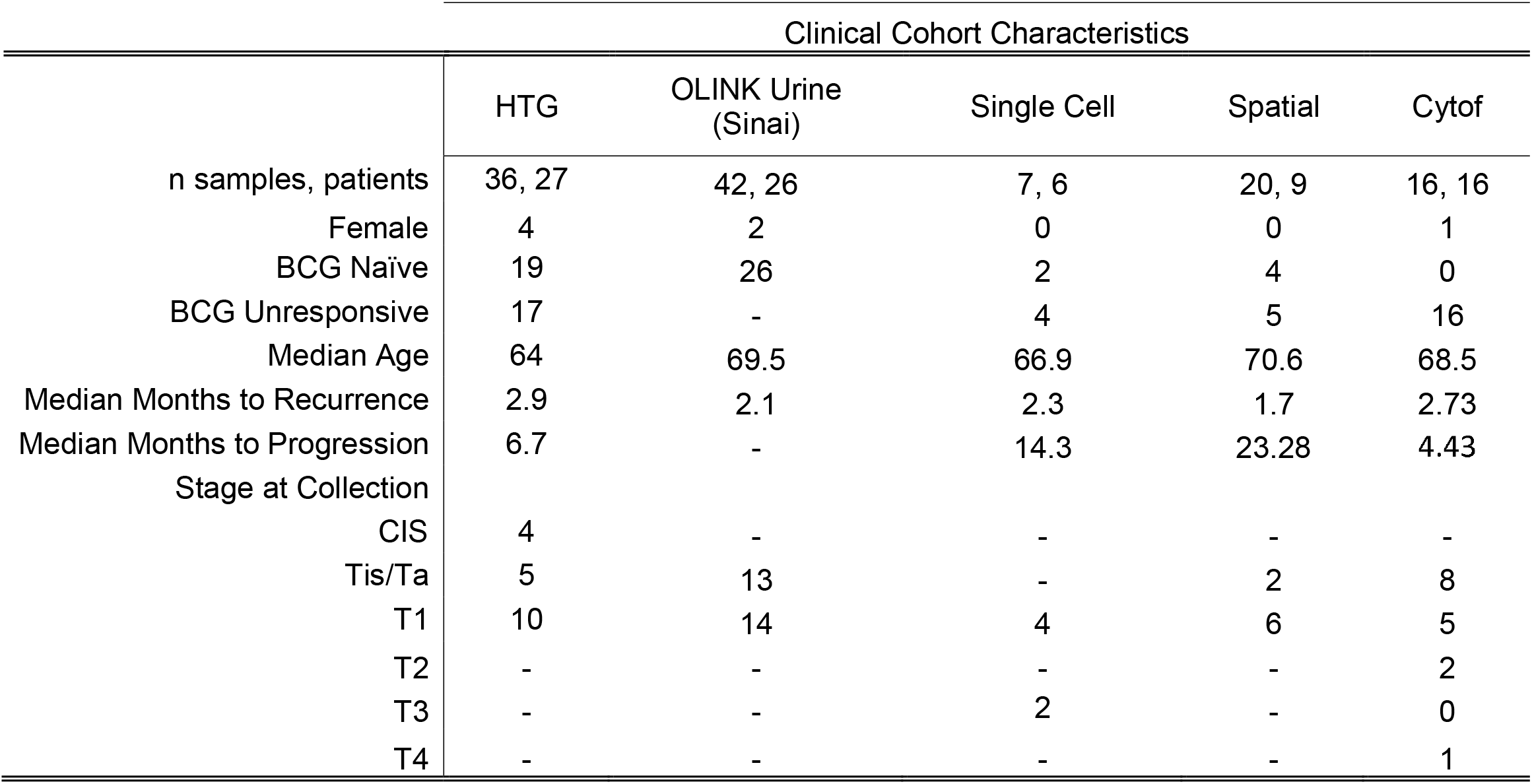
Clinical characteristics of experimental cohorts.

### IFN-γ mediates adaptive resistance in BCG-unresponsive tumors

To investigate a potential mechanism of adaptive resistance in NMIBC, we used tumor specimens from BCG-unresponsive and BCG naïve tumors collected from a longitudinal cohort of NMIBC patients. Targeted *in situ mRNA* sequencing was performed, and a gene-set enrichment analysis (GSEA) comparing BCG-unresponsive tumors to pre-treatment timepoints was run (**Fig. 1A**) (24). Statistically significant gene sets are shown ranked by their normalized enrichment score (NES) (**Fig. 1A**, **Supplemental Table S1**). The top enriched gene-set comparing BCG-unresponsive to BCG naïve tumors (n=36) includes IFN-γ signature, B cell markers, cytotoxic lymphocytes, lymphocyte receptors/function, T cell checkpoint modulation, tumor inflammation, TH1 response, lymphocyte trafficking, general T cell markers, and receptor genes (**Supplemental Table S1**). Gene sets that are downregulated in the BCG-unresponsive state and upregulated in the BCG naïve state are TGF beta signaling, hypoxia, peroxisome, oxidative phosphorylation, P53 pathway, DNA repair, and stress toxicity.

**Figure 1:**
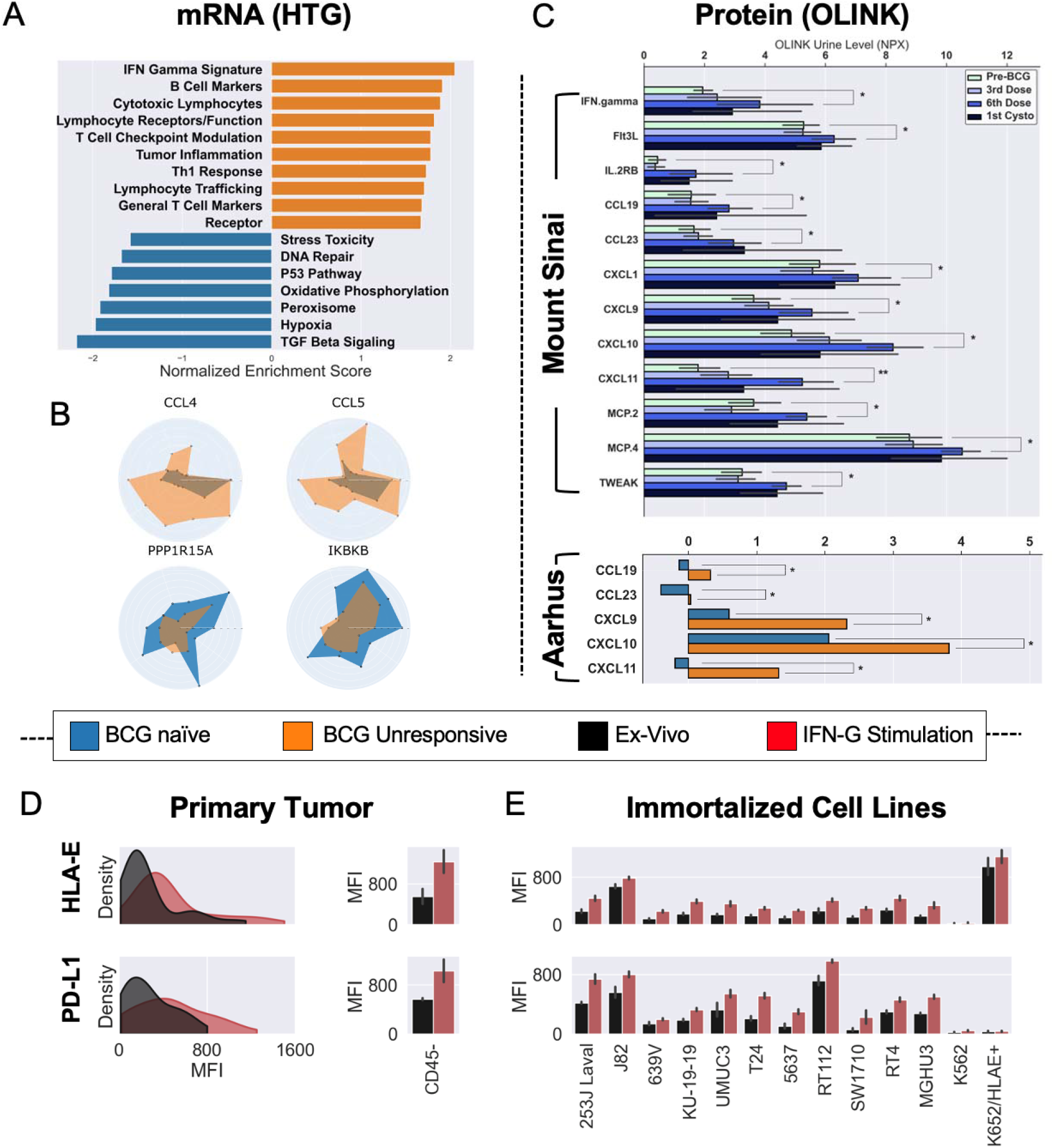
Adaptive resistance in BCG unresponsive disease revealed by targeted RNA sequencing and OLINK proteomics. **A:** Targeted RNA gene-set enrichment analysis showing statistically significant differences between BCG naïve and BCG unresponsive cases. IFN-γ signature is the most highly upregulated signature in the BCG unresponsive patients. All gene sets are significant at p < 0.05 by Kruskal-Wallis or independent T-test. **B:** Radar plots, one patient per spoke, showing upregulated individual genes (*CCL4* and *CCL5*), and downregulated individual genes (*PPP1R15A* and *IKBKB*). All genes significant at by Kruskal-Wallis or independent T test (p < 0.05). **C:** Longitudinal protein analysis from Mount Sinai in all patients (N=27) with th comparisons between BCG naïve and 6^th^ induction dose timepoints (* p < 0.05, ** p < 0.001). Validation cohort (n=66) from Aarhus showing BCG naïve vs BCG unresponsive timepoints in urine samples (p values in both cohorts assessed via independent T test or Kruskal-Wallis with Benjamini-Hochberg correction for multiple comparisons). **D:** MFI of primary non-muscle invasive bladder cancer cells after 24 hours of incubation with IFN-γ. **E:** MFI of commercial immortalized cell lines after 24 hours of IFN-γ stimulation (all values p < 0.05). All MFI experiments performed in triplicate.

Differences in individual genes between paired BCG naive and BCG-unresponsive samples reveal that all paired samples show upregulation of CCL4 and CCL5 and near-universal downregulation of PPP1R15A and IKBKB (**Fig. 1B**; all p < 0.05). CCL4 and CCL5 are potent chemoattractants for a variety of cells, among which are NK and T cells (25,26). IKBKB is an activator of NF-κB-dependent apoptotic pathways, and PPP1R15A has functions associated with negative regulation of TGF-b signaling (27,28).

Validating our *mRNA* analysis at the protein level, we used the OLINK^TM^ “inflammation panel,” a 92-analyte protein proximity extension assay, to profile urine cytokines reflective of the urothelial microenvironment (29,30), from our longitudinal BCG-treated cohort of NMBIC patients. Upregulated cytokines of interest in the sixth dose compared with the BCG naïve state include IFN-γ, Flt3L, IL2RB, CCL8, CCL13, CCL19, CCL23, CXCL1, CXCL9, CXCL10, CXCL11, and TWEAK (**Fig. 1C**; all p<0.05 following Benjamini-Hochburg correction).

The inflammatory signature seen during the induction sequence at Mount Sinai was validated in a longer-term OLINK cohort from Aarhus University. Urine samples from 66 patients were collected at the BCG naïve time point and compared to urine samples from the same patients at the time of recurrence. CXCL9, CXCL10, CXCL11, CCL19, and CCL23 had lasting upregulation in the BCG unresponsive timepoint (p < 0.05 following Benjamini-Hochburg correction).

To better understand the effects of IFN-γ on tumor dysregulation, we incubated primary tumor cells from NMIBC patients (n=10) and commercial urothelial cell lines in the presence of recombinant human IFN-γ (**Fig. 1D** and **E**) and measured HLA-E and PD-L1 via fluorescence flow cytometry after 24-hours in triplicate experiments. A histogram (left) and bar plot (right) show sorted CD45-cells from primary NMIBC tumors before and after IFN-γ stimulation (**Fig. 1D**). The CD45^-^ cells show statistically significant increases in PD-L1 and HLA-E following stimulation with IFN-γ. Expanding the analysis to commercial immortalized cell lines (n=11) shows the signature is consistent: PD-L1 and HLA-E experienced a statistically significant increase after 24-hour incubation with IFN-γ **(Fig. 1E)**. Further, we saw no effects on HLA-E expression in WT K562 cells or HLA-E–transfected K562 cells, in which HLA-E expression is controlled under a very strong CMV promoter.

### NK and CD8 T cell-derived IFN-□ drives HLA-E expression on BCG-unresponsive tumors through dysregulated STAT1

Single-cell RNA sequencing (scRNA-seq) on five NMIBC samples, BCG naïve and unresponsive, was performed to refine the associated IFN-γ genes further (**Supplementary Table S2**). 24,166 cells containing 26,520 gene features passed quality control and were used for subsequent analyses, with a median of 958 genes detected per cell. (**Supplementary Fig. S1**).

Twenty-eight identifiable cell populations, including CD8^high^ T cells, regulatory T cells (Tregs), non-CD8 (*CD4^-^ CD8A^-^ FOXP3^-^*) CD8^low^ T cells, NK cells, dendritic cells (DCs), monocytes and macrophages, granulocytes, B cells, plasma cells, and nonhematopoietic fibroblasts, epithelium, and tumor cells were identified after graph-based clustering (**Fig. 2A**, **Supplementary Table S3**). To identify the cell origins of IFN-γ and related pro-inflammatory gene signatures, we examined the expression of *IFNG* in each of the cell populations. We found that tissue-resident memory (TRM) CDs^high^ T cells, canonical CD8^high^ T cells, NK cells, and CD8^low^ T cells demonstrated the highest level of *IFNG* expression among all cell clusters and have distinct IFN-γ^high^ cell populations (**Supplementary Fig. S2).**

**Figure 2:**
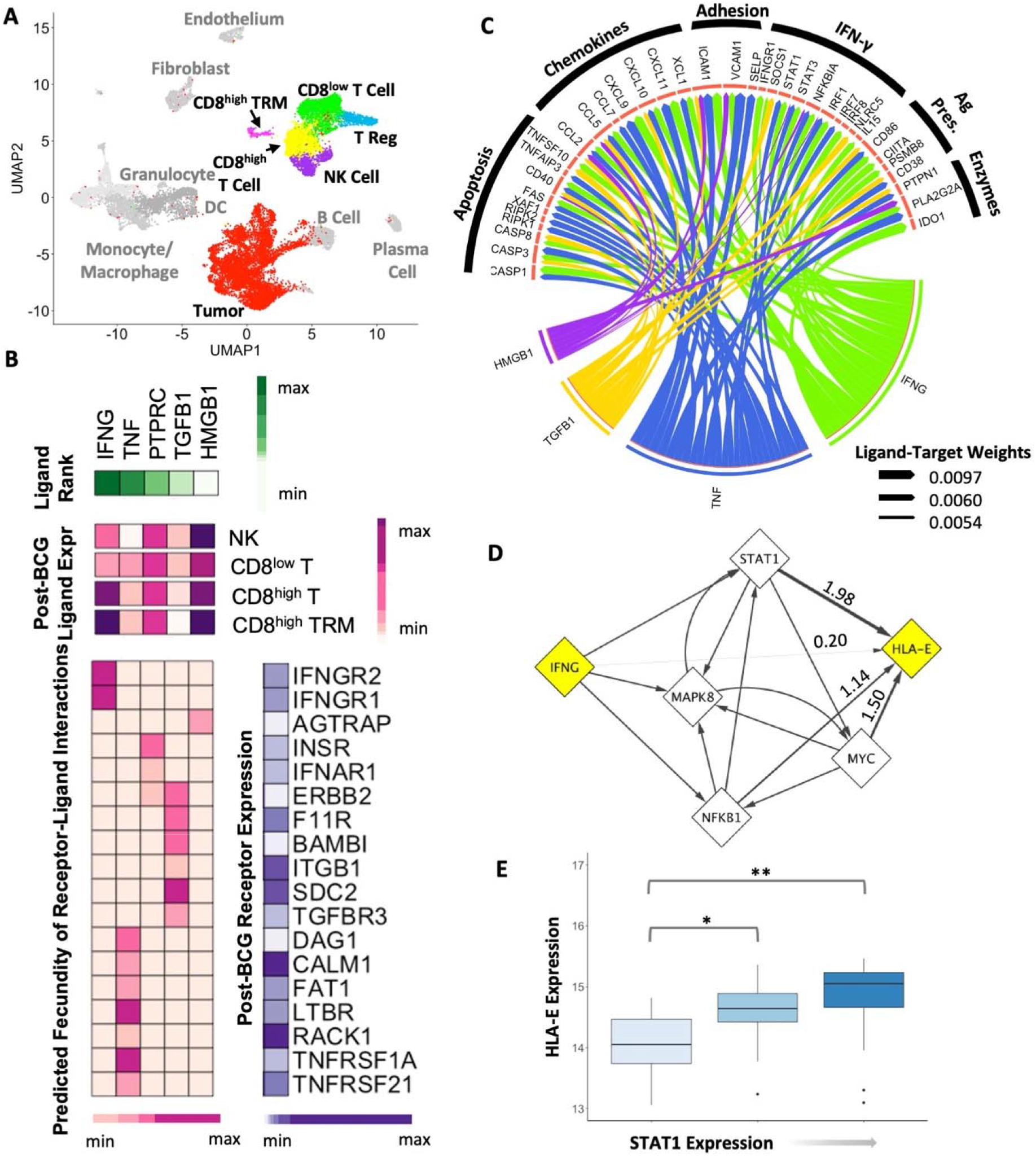
IFN-γ signaling is predicted to drive downstream HLA-E expression via STAT1 in tumor cells. **A:** UMAP visualization of cell types from scRNA-Seq analysis of urothelial tumor samples. **B:** NicheNet predictions of upstream ligand-receptor interactions showing *IFNG, TNF, PTPRC, TGFB1*, and *HMGB1* (left) as the top 5 ligands expressed in NK, CD8^LOW^ T, CD8+ T, and CD8+ TRM cells (middle) that most likely influence tumor cell expression of interferon-gamma signaling genes. Strength of ligand-receptor interactions (top right) and expression of the receptors in tumor cells (bottom right) are also shown. **C:** Circos plot showing links between NicheNet-predicted ligands and downstream interferon-gamma pathway genes. Strong relationships between ligand-target gene pairs are demonstrated by both wider arrow width and solid arrow colors. **D:** NichetNet-predicted regulatory network between IFN-γ and HLA-E. **E:** Expression of HLA-E in HTG targeted RNA-Seq urothelial cancer cohort with respect to STAT1 levels. (*p = 0.0274; **p = 0.00674)

Although IFN-γ can have anti-tumor properties via activation of cellular immunity, it has also been shown to be involved in tumor growth. To study the response to IFN-γ, we investigated the effects of NK and T cell-derived IFN-γ signaling on tumor cells. To do so, we employed NicheNet, an algorithm that uses a combination of transcriptomic data, known ligand-receptor interactions, and gene-gene signaling and regulatory relationships to predict potential interactions between cells of interest (31). The top five ligands from CD8^high^ T cells and NK cells predicted to induce tumor expression of IFN-γ response genes were interferon-gamma (*IFNG*), tumor necrosis factor (*TNF*), CD45 (*PTPRC*), transforming growth factor (TGF)-beta (*TGFB1*), and high mobility group box 1 (*HMGB1*) (**Fig. 2B**). TRM CD8^+^ T cells expressed the highest levels of *IFNG* and *HMGB1* relative to NK and CD8^+^ T cells, while *TNF* and *TGFB1* expressions were highest in non-TRM CD8^+^ T cells and NK cells, respectively (**Fig. 2B**). Putative receptors of these ligands, including *IFNGR1, SDC2*, and *TGFBR3*, were more highly expressed in the BCG-unresponsive samples, suggesting the presence of corresponding signaling activity in the tumor (**Fig. 2B**). To further examine genes upregulated in the tumor as a consequence of signals from NK and T cells, we created a circos plot demonstrating the ligand-target weights of the top third of predicted relationships between ‘sender’ or effector ligands and ‘receiver’ target genes (**Fig. 2C**). Aside from CD45, all other top-ranked ligands were predicted to induce expression of IFN-γ response genes to varying degrees. The strongest predicted target of *IFNG* and *TNF* was intercellular adhesion molecule 1 (*ICAM1*), a cell surface glycoprotein commonly found expressed on endothelial cells and immune cells with roles in cell migration. Additionally, genes involved in apoptosis, TNF-related ligands and receptors, inflammatory response, and interferon-gamma signaling were also predicted to be upregulated as a result of sender ligand signaling by tumor cells (**Fig. 2C**).

A downstream target of IFN-γ signaling, *HLA-E* is predicted to be under direct regulation of transcriptional regulators *STAT1, MYC*, and *NFKB1*, with *STAT1* predicted to have the strongest regulatory potential for *HLA-E* expression (**Fig. 2D**, **Supplementary Table S3**). Among these, *STAT1* is also a target gene that is upregulated in bladder tumor cells as a result of IFN-γ signaling from NK and CD8^+^ T/TRM cells (**Fig. 2C**). We further confirmed the *STAT1-HLA-E* axis in the in situ targeted *mRNA* analysis of NMIBC patients, which demonstrated a direct positive correlation (Spearman rho = 0.455, p-value = 2.74e^-3^) between *STAT1* and *HLA-E* expression (**Fig. 2E**). Though *MYC, NFKB1*, and *IFNG* were also predicted to exert some degree of regulatory effect on *HLA-E* expression, these genes failed to show a direct correlation with *HLA-E* expression by targeted *mRNA* analysis (**Supplementary Fig. S2**).

### NK cells, CD8 T cells, and Tregs preferentially infiltrate activated tumor “nano-environments” with higher densities of HLA-E and CXCL9/10/11

To spatially resolve the relationship between tumor nests and immune infiltrates, we performed a series of 10X ST-seq analyses. Twenty total slices of tumor were sequenced from four treatment-naïve and four BCG-unresponsive tumors, including one unresponsive tumor treated with BCG and PD-1 blockade (**Fig. 3**). Spatial relationships were investigated on the ‘nano-environment’ level, defined as a Visium dot with a radius of 2.5 adjacent Visium spots, or roughly ~138 microns. A representative BCG unresponsive tumor section with a topographical map of HLA-E expressing cell density is shown on top of a neighbor’s analysis highlighting the intensity of neighboring CD4^+^ T cell infiltration (**Fig. 3A**) and NK and CD8^+^ cell infiltration (**Fig. 3C**). An alternative view of binarized HLA-E density with CD4^+^ T cells in red (Fig. 3B) and NK/CD8^+^ T cells in magenta and red, respectively (**Fig. 3D**), is also presented. *HLA-E*-expressing (*HLA-E^HIGH^*) Visium spots are shown in blue, and no/low *HLA-E* Visium spots are shown in tan. We observed a very clear clustering of CD8 T cells around HLA-E^HIGH^ spots and high- *HLA-E* density areas. Conversely, we observed only sparse NK or T cells overall in the treatment-naïve tumor sections (**Supplemental Fig. S3**).

**Figure 3:**
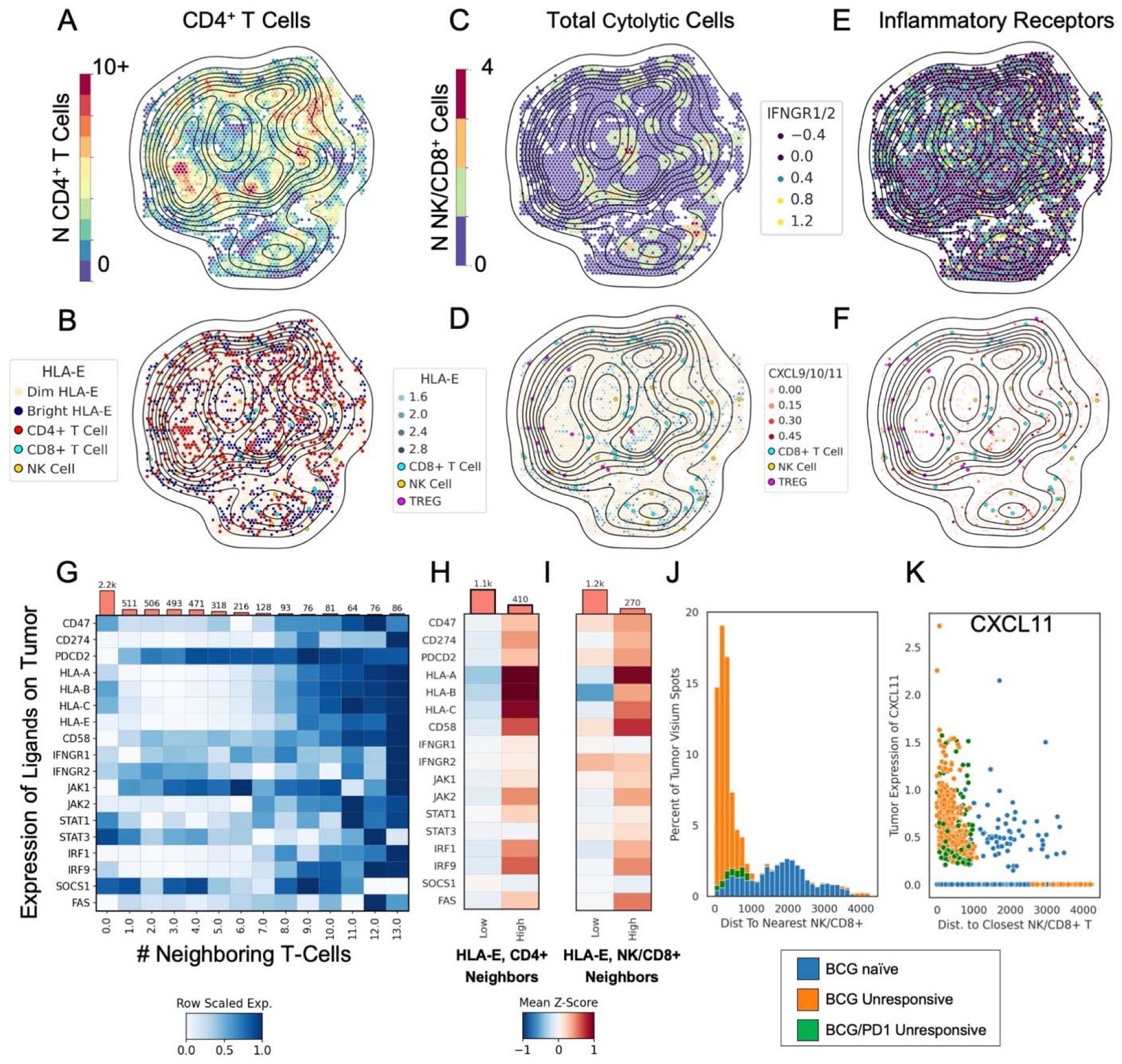
NK cells, CD8 T cells, and Tregs preferentially infiltrate activated tumor “nano-environments” with higher densities of HLA-E and CXCL9/10/11. **A:** representative spatial slice colored by number of neighboring infiltrating CD4^+^ T cells, with a topographical plot showing density of *HLA-E* expression. **B:** same representative slice showing all T cells, CD8^+^ cells, and NK cells with *HLA-E* expression shown in topographical plots, as well as low and high expressing cells in tan and blue. **C:** representative spatial slice colored by number of neighboring infiltrating cytotoxic cells (CD8+ and NK cells), with a topographical plot showing density of HLA-E expression. **D:** NK and CD8+ T cells next to *HLA-E^HIGH^* cells. **E:** representative spatial slice colored by *IFNGR1* and *IFNGR2* score with a topographical plot overlaid showing density of *HLA-E* expression. **F:** representative spatial slice showing *CXCL9/10/11* score, T cells, NK cells, and an overlaid topographical plot of *HLA-E* density. **G:** meta-analysis of all tumor-labelled spatial points showing row-scaled expression of pertinent genes stratified by number of neighboring infiltrating CD4^+^ T-cells. **H:** Z-scored heatmap showing *HLA-E^LOW^*, low-CD4^+^ infiltrating tumor spots (n = 1.1K) vs *HLA-E^HIGH^* and high-CD4^+^ infiltrated tumor spots (n=410). **I:** Z-scored heatmap showing HLA-E^LOW^, low-cytotoxic infiltrating tumor spots (n = 1.2K) vs HLA-E^HIGH^ and high-cytotoxic infiltrated tumor spots (n=242). **J:** histogram showing the percent of tumor-expressing Visium spots by distance to nearest NK or CD8^+^ cell. **K**: *CXCL11* expression on tumor-identified Visium spots as a function of the distance to the closest neighboring NK or CD8+ T cell.

In order to investigate the effects of inflammatory signaling, we profiled the same representative tumor slice for IFN-□ and downstream cytokine signaling. First, *IFNGR1/2* expression was scored and plotted alongside *HLA-E* density (**Fig. 3E**). Strikingly we found the greatest expression of *IFNGR1/R2* in areas of dense *HLA-E* expressing cells. Second, in order to understand potential mechanisms of preferential infiltration of *HLA-E^HIGH^* tumor nests, we profiled chemokine expression by tumors. Interestingly, we observed that *HLA-E^HIGH^* tumors were significantly more enriched for elevated expression of *CXCL9/ CXCL10/CXCL11* (**Fig. 3F**). Collectively, the data demonstrate that *HLA-E^HIGH^* tumors are activated to recruit T cells. Importantly, this observation was unique to *HIA-I^HIGH^* tumors and was not seen in treatment-naïve specimens (**Supplemental Fig. S3**). These data are supported by two previous studies of melanoma showing that IFNs (type I or II) induce CXCL9/10/11 by tumors and attract CXCR3^+^ T cells to the TME.(32,33)

Expanding to a meta-analysis of the entire cohort of tumor sections, we performed a differential gene expression analysis on Visium spots identified as encompassing tumor cells stratified by the number of neighboring infiltrating CD4^+^ T cells (**Fig. 3G-3K)**. We observed significantly greater numbers of CD8^+^ T cells surrounding *HLA-E^HIGH^* tumor spots (**Supplemental Fig. S4**). We next profiled the expression of key inhibitory ligands expressed on tumors *CD47, CD274* (PD-L1), *PDCD2* (programmed cell death protein 2), and *CD58*, along with *HLA-A, -B, -C, -E*, and *IFNGR1/R2* (**Fig. 3G**). Expression levels increased on tumor spots as the numbers of infiltrating CD4^+^ T cells increased (p < 0.05 following correction for multiple hypotheses). The *class I HLA* genes were most sensitive to increasing T cell infiltration, followed by *CD58* and *CD274* (PD-L1). Additionally, interferon pathway genes were also enriched in the tumor nests that are more intensely infiltrated by T cells: *JAK1, JAK2, STAT1, IRF1, IRF9*, and *FAS* were all significantly upregulated. <u>Importantly, *SOCS1* and *STAT3* were not upregulated with increasing tumor infiltration</u>. Whereas STAT3 signals are critical for the production of IL-6 and IL-10, Suppressor of Cytokine Signaling Protein-1 (SOCS1) plays a critical role in negative feedback inhibition to shut down inflammatory immune responses through targeting JAK1 and JAK2 for proteasomal degradation (34–36).

Finally, the spatial neighbors’ analysis was expanded to investigate Visium spots containing tumors stratified by HLA-E and infiltrating T cells, with high infiltration being >= seven neighboring T cells and low infiltration being <= four neighboring T cells. The *HLA-E^LOW^* condition had only 90 tumor Visium spots, or 6.5%, classified as being highly by infiltrated T cells; the *HLA-E^HIGH^* condition had 410, or 29.8%, of tumor Visium spots classified as being highly infiltrated by T cells. Further, the *HLA-E^HIGH^* tumor Visium spots showed the same upregulation as when classified by highly infiltrated by tumor cells and were enriched in *CD47, CD274* (PD-L1), *PDCD2, HLA-A, -B, -C*, -*E*, *CD58, IFNGR1, IFNGR2, JAK1, JAK2, STAT1, IRF1, IRF9*, and FAS (**Fig. 3H, 3I)**.

Lastly, in order to reinforce the spatial relationship between cytolytic cells and tumor-specific cytokines, we calculated the distance on each slice between every Visium spot and their nearest neighboring cytolytic cell (NK^+^ or CD8^+^ T cell). BCG unresponsive and BCG / PD-1 unresponsive tumor samples make up the vast majority of tumor spots in close proximity to cytolytic cells (**Fig. 3J).** When plotted as a function of distance to nearest cytolytic cell (NK or CD8^+^), tumor expression of *CXCL11* increases as proximity to cytolytic cells increases (**Fig 3K).** Notably, this phenomenon exists almost exclusively in the BCG unresponsive state. *CXCL11* expression is ~55 fold higher in the BCG unresponsive state when compared to the BCG naïve state (p < 0.001). Additionally, BCG unresponsive tumors see statistically significantly higher levels of infiltration with cytotoxic and regulatory T cells: Tregs are a median of 2.66 times closer, and cytolytic cells are a median of 7.05 times closer to tumor-identified Visium spots in the BCG unresponsive state when compared to the BCG naïve state (p < 0.0001 for both; **Fig. 3J, K; Supp Fig S5**).

### *KLRC1* (NKG2A) expression stratifies NK and CD8 T cell capacity for anti-tumor functions in BCG-unresponsive tumors

Given the hypothesis that NKG2A presents a novel immune checkpoint receptor in bladder cancer, we sought to characterize the cytolytic capacity of immune populations most impacted by NKG2A expression. Stratifying by *KLRC1* (NKG2A) expression on immune populations defines distinct phenotypes of cytolytic activity. (**Fig. 4A**). Our data has identified that *KLRC1^high^* CD8 T cells expressed elevated *GNLY, TIGIT, GZMA, GZMB, HAVCR2, LAG3*, and *CTLA4*. CD8 TRM cells showed significant upregulation in *GNLY, GZMA, GZMB, LAG3, TIGIT*, and *HAVCR2* in the *KLRC1* cells. Finally, a similar analysis of NK cells showed significant upregulation in *XCL1, TIGIT, XCL2*, and *GZMK* between *KLRC1* cells.

**Figure 4:**
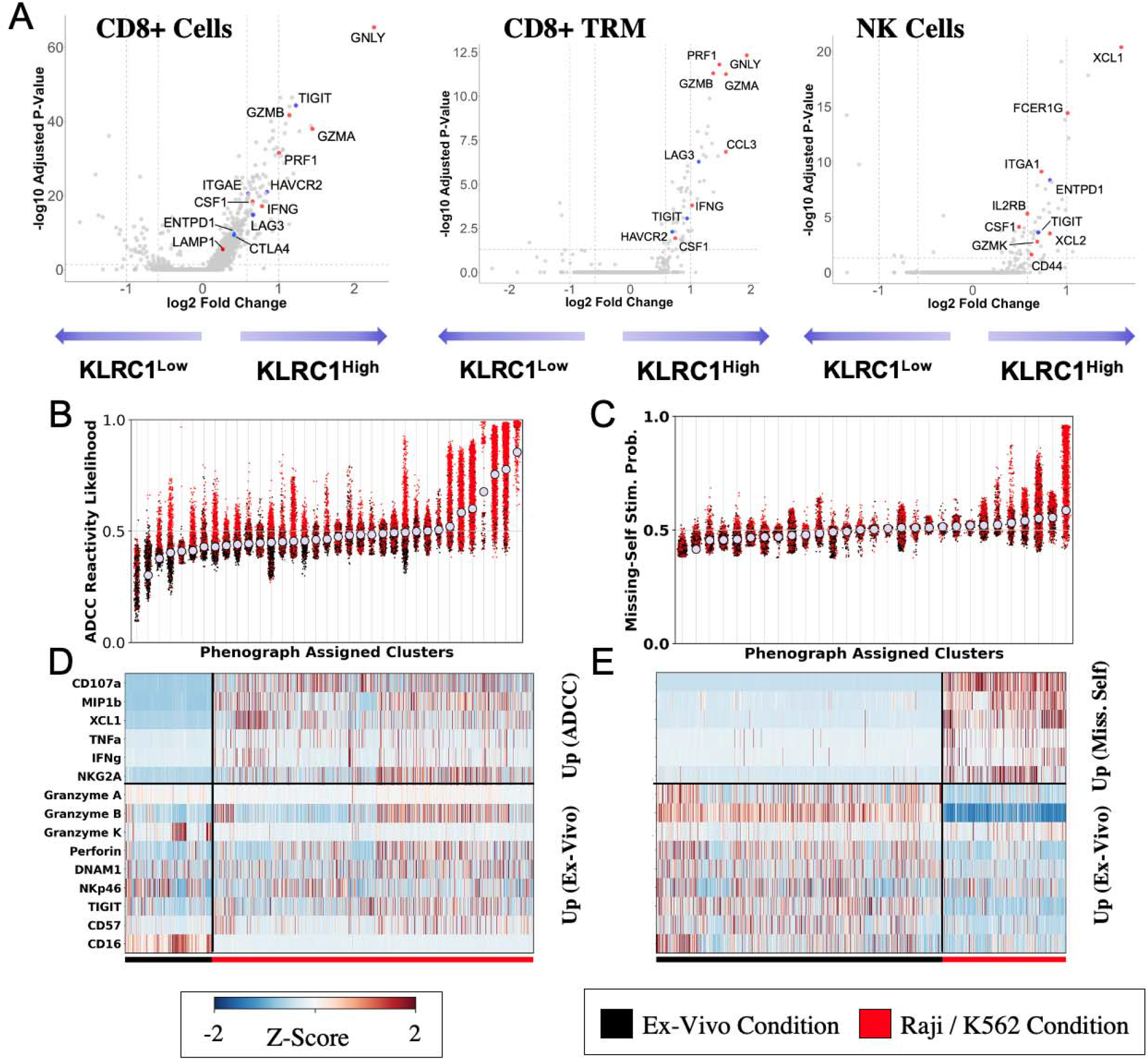
*KLRC1* (NKG2A) expression stratifies NK and CD8+ T cell capacity for anti-tumor functions in BCG unresponsive tumors. A: single-cell RNA sequencing showing statistically significant upregulation of both immune checkpoints and cytotoxic genes in KLRC1+ CD8+ T cells, CD8+ TRMs, and NK cells. B: phenograph clusters stratified by ADCC reactivity. C: phenograph clusters stratified by missing-self reactivity. D: CYTOF markers partitioned by strongly ex-vivo vs strongly ADCC reactive clusters. E: CYTOF markers partitioned by strongly ex-vivo and strongly missing-self-reactive clusters.

### Blood-derived NK cells retain function competence at time of tumor recurrence

To better understand if an elevated expression of cytolytic genes in NKG2A^+^ NK, CD8 T, and TRM cell populations correlated with improved functional sensitivity, we used mass cytometry (CyTOF) to profile the *in vitro* response to BCG by blood-derived NK cells from NMIBC patients at the time of tumor recurrence. We performed a phenograph clustering analysis (37) to stratify the data by their likelihood of existing in the ex-vivo state vs. the experimental state, using either an assay of antibody-dependent cellular cytotoxicity (ADCC) reactivity or loss of class I HLA ‘missing-self’ stimulation conditions. Highly polarized clusters of immune cells underwent a differential protein expression analysis, MELD, comparing levels of important markers that dominantly defined the ex-vivo vs. highly experimental conditions (38).

Blood-derived NK cells at the time of recurrence experience retained anti-tumor abilities when stimulated by ADCC-inducing or missing-self conditions (**Fig. 4B** and **C**). Clusters highly likely to be found in the ADCC-induced condition after 6 hours of stimulation express statistically significantly higher levels CD107a, MIP1b, XCL1, TNFα, IFN-γ, and NKG2A. The ADCC-induced condition expressed statistically significantly lower granzymes a and k, NKp46, and CD16 suggesting tumor lysis as well as maturation during the time of co-culture. Additionally, activating receptors, which are known to be down-regulated during maturation (NKG2D, NKp46) were also downregulated. The terminal exhaustion maturation marker, CD57, was upregulated as well, suggesting the dominantly activated NK cells were maturing and active, but also expressing exhaustion markers. NK cells highly polarized towards the missing-self condition after 6 hours of stimulation and showed statistically significant increases in expression of CD107a, MIP1b, XCL1, TNFα, CD56, IFN-γ, and NKG2A; and statistically significant decreases in granzymes a and k; perforin; DNAM1; NKp46; TIGIT; CD57; and CD16. This suggests a slightly different picture, where exhaustion marker CD57 remains low, indicating that the NK cells are maturing and active without similarly induced exhaustion as the ADCC condition (**Fig. 4D** and **E**).

## Discussion

Adaptive immune resistance can be driven by tumor-intrinsic and/or extrinsic mechanisms in overcoming cytolytic immune clearance (39,40). Our group recently demonstrated that anti-tumor adaptive immune responses are often observed along with a pro-tumorigenic inflammatory response, but the balance between these competing signatures (defined as the 2IR score) predicts response to anti-PD-1/PD-L1 blockade in metastatic bladder cancer patients treated with (40). Evidence from mouse and human studies demonstrates that interferons, while stimulating a robust anti-tumor response, also upregulate immune-suppressive factors in the setting of prolonged activation. In melanoma, IFN-γ from CD8 T cells was shown to upregulate PD-L1 and mediate enrichment of FOXP3^+^ regulatory pathways within the TME (41). In a broader meta-analysis across 18 tumor indications, including bladder cancer, inflammatory mediators (e.g., IFN-γ) were associated with inhibitory immune checkpoints, including PD-L1/L2 (42,43). Despite the emerging evidence positioning pro-tumorigenic roles for IFN-γ, there are well-established anti-tumor functions mediated through IFN-γ that are critical for anti-PD-L1 immunotherapy (44). Anti-tumor inflammation, therefore, exists along a continuum where an equilibrium is necessary for appropriate immunotherapeutic efficacy (**Fig. 5**) (45).

**Figure 5:**
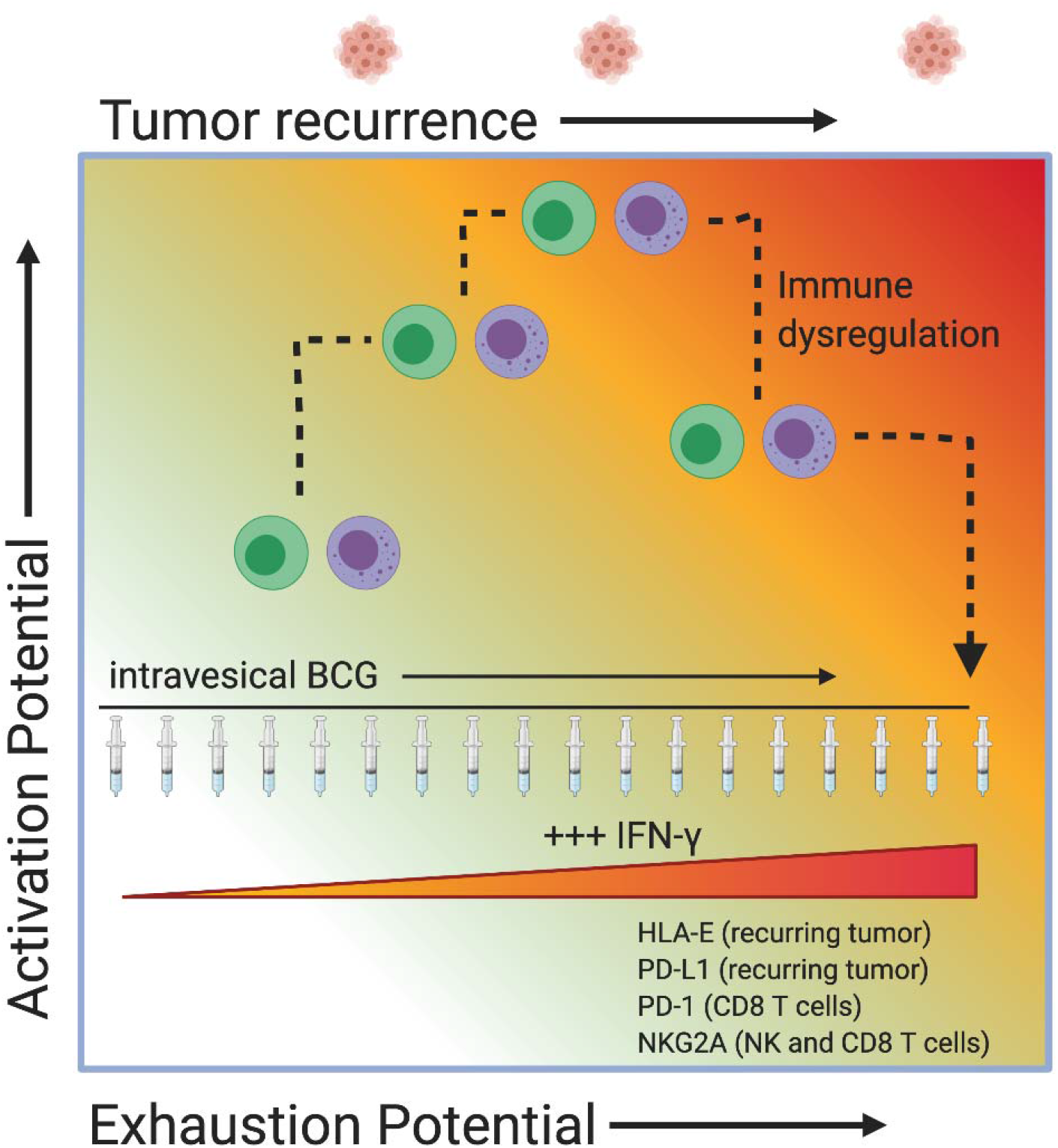
Schematic of proposed mechanism of immune dysregulation in the setting of chronic immune activation via BCG administration.

In this study, we investigated the impact of BCG-induced inflammation on the interplay between tumor and immune cell populations, using both prospective and retrospective NMIBC specimens before and after immunotherapy. Our data suggest: i) inflammation is ubiquitous in BCG-responders and non-responders alike; ii) IFN-γ directly contributes to the upregulation of HLA-E and PD-L1 on recurring NMIBC tumors; iii) tumors stratified by neighboring infiltrated NK and T cells reveal a program of activation (and possibly dysregulation) alongside increased immunosuppression possibly mediated by Tregs; vi) unique to BCG-unresponsive tumors, HLA-E^HIGH^ tumor nests were strongly activated to produce CXCL9/10/11, which correlates with infiltration by NK and CD8 T cells (and Tregs); v) NKG2A expression stratifies NK and CD8 T cell anti-tumor cytolytic functions, however, NKG2A-expressing NK and CD8 T cells also expressed a number of well-established secondary checkpoint receptors; vi) finally, NK cells removed from their suppressive microenvironment demonstrated retained functional capacity. These findings frame a picture of immune dysregulation dominantly defined by elevated tumor expression of HLA-E and PD-L1 in settings of BCG-induced chronic activation and inflammation.

Intravesical administration of BCG is initiated as adjuvant immunotherapy, as complete tumor resection is the initial diagnostic and therapeutic step. Given the context, the current standard of care (SOC) BCG immunotherapy presupposes inflammation without active tumor to clear (46). The development of improved treatments for NMIBC and BCG-resistant disease has lagged. This may be, in part, because few studies have attempted to understand the relationship between the timing of tumor recurrence, reasoning for the recurrence, and the state of the immune system at the time of recurrence. Poor dosing study designs and lack of understanding of the mechanisms underlying a therapeutic response to intravesical BCG have led to a significant gap in knowledge and treatment improvements for patients with NMIBC compared to muscle-invasive or metastatic disease.

Our findings suggest that all NMIBC patients at the time of tumor recurrence show signs of a hallmark anti-tumor immune response dominantly driven by IFN-γ. BCG-unresponsive tumors see uniform increases in chemotactic cytokines and inflammatory pathways that should otherwise function to suppress their growth. Further, increased expression of inhibitory ligands on BCG-unresponsive tumor cells was observed, suggesting that inflammatory stimuli had been prolonged and triggered feedback mechanisms responsible for immune evasion. Thus, when tumors recur, for reasons beyond the scope of this study, they are met by a functional status of the immune system ill-equipped to suppress them.

Previous analyses profiling urine analytes between the BCG naïve and third dose timepoints demonstrated that three doses of BCG induced an inflammatory response hundreds of times above baseline levels. IP-10 was found to be 227.5 times increased over baseline levels alongside MIP-1ß (82.6 fold increase), IL-8 (56.7 fold increase), IL-6 (44.4 fold increase), and TNFα (11.4 fold increase), among others (47). Importantly, the third dose-response dwarfed the magnitude of the first dose-response, suggesting that repeated exposure increases the magnitude of inflammation (47). While these data did not profile out to the time of tumor recurrence, they lend credence to the theory that all patients experience a ubiquitous and increasingly powerful immune response to repeated doses of BCG.

IFN-γ is well established as both a response to BCG stimulation and as a mediator of elevated HLA-E expression via JAK/STAT pathways, which are shared with other HLA class I genes (48–51). In our neural network analysis (**Fig. 2**), IFN-γ enhanced expression of HLA class I, including HLA-E, via JAK-1 and JAK-2 and STAT1 pathways, but was uniquely restricted to BCG-unresponsive tumors suggesting these tumors were dysregulated. In fact, we observed direct effects of IFN-γ on increasing HLA-E expression across all analyses. Given our data on the correlation between STAT1 tertials and *HLA-E* expression, and supported by mechanistic studies on IFN-γ signaling, our data offer a potential mechanism of dysregulated IFN-γ signaling in BCG-induced inflammation, which directly leads to elevated HLA-E expression on recurring tumors.

Novel to our study are the spatially-resolved insights into potential consequences of IFN-γ dysregulation on immune homing patterns. Preferential infiltration of HLA-E^HIGH^ and PD-L1^HIGH^ tumor nests by immune cells was seen in areas of high tumor production of CXCL9/10/11 (**Fig. 3**). Further, BCG unresponsive tumors saw a more than 50-fold increase in production of CXCL11 when compared to BCG naïve nests, and tumor proximity to cytolytic cells and Tregs was increased a median of 7-fold and 2.6-fold respectively. The phenomenon of tumors producing CXCL9/10/11 has been initially observed in melanoma, where IFN-γ induced CXCL9/10/11 production, increasing the chemotactic gradient for CXCR3^+^ T cells (33,52). In our data, CXCL9/10/11-producing tumor nests show upregulated mediators of activation, including *IRF1, IRF9*, and *JAK2*, as well as absent expression of cytokine-suppressing mediators, like *SOCS1* (9). These nests are notably upregulated for immunosuppressive ligands as well: highly T and NK cell-infiltrated tumor areas express higher levels of *HLA-E, CD47, CD274* (PD-L1), and *PDCD2* (programmed cell death protein 2). To the best of our knowledge, this is the first demonstration where spatial resolution of cell sequencing shows that tumor-derived CXCL9/10/11 chemotactically recruits NK and T cells (including regulatory T cells) as a mechanism of sparing HLA-E^LOW^ PD-L1^LOW^ tumors that might otherwise be susceptible to lysis.

Clinical efficacy in checkpoint blockade is dependent on reinvigorating effector cells expressing high levels of the targeted checkpoint(s). Recent studies have demonstrated that expansion and functional sensitivity of circulating CD8 T cells predicts response to PD-L1 blockade and mechanistically provide a pool of replenishing effectors that traffic to the TME (53). Congruent with analogous studies, we found that i) *KLRC1*^HIGH^ NK and CD8 T cells were enriched in BCG-unresponsive tumors, ii) circulating NK cell anti-tumor functions, defined by ADCC and missing-self reactivity, were intact at the time of tumor recurrence, and iii) the NKG2A^HIGH^ expressing NK and CD8 T cells reflected the dominantly activated phenotype (**Fig. 4**). Previous studies have demonstrated a similar phenomenon, with PD1^HIGH^ TIGIT^HIGH^ CD8 T cells in melanoma expressing high levels of IFN-γ, GZMK, and IL10, alongside higher *TOX* levels known to be a marker of T cell exhaustion (54,55). Additionally, these cells expressed variable CXCR3 levels with only peripheral CD8^+^ T cells expressing CXCR3 (55). Peripheral blood-derived NK and T cells have become increasingly important as clonal expansion of T cells predictive of immunotherapeutic response in several tumor phenotypes have been seen to be reflected clearly in the periphery (53,55,56). This reinforces both the use of NKG2A to stratify cytolytic effector genes in NK, CD8 T, and CD8 TRMs, as well as the notion that CXCL9/10/11 lures in effector cells to take on exhausted phenotypes.

This study has notable limitations. Our sample sizes, while being the largest NMIBC spatial sequencing cohort available, are small and present a potentially limited view of the disease. Further, both spatial sequencing and single-cell RNA sequencing have known difficulties detecting transcripts from a large proportion of transcribed genes, several of which were pertinent to our study. Spatial sequencing poorly transcribed *CD4, CXCR3, FASL, GAS2, GAS3, IFN-γ*, and *KLRC1* were poorly expressed in our spatial dataset.

Collectively, our analyses suggest that the current guidelines on immunotherapy in NMIBC could be improved via a multi-cell-targeting immunotherapy approach focused on NK cells alongside T cells. Randomized trials from bladder and other tumor indications have shown that PD-1/PD-L1 stratification fails to predict response to immunotherapy with anti-PD-1/PD-L1 antibodies: IMvigor210, JAVELIN bladder 100, and CheckMate-275 (NCT02108652, NCT01772004, NCT02387996) all saw that PD-1/PD-L1 biomarker stratification alone did not effectively stratify response rates. In comparison, recent results using *KLRC1* (NKG2A) expression in the pre-treatment tumor successfully predicted anti-PD-L1 response rates in the IMvigor210 cohort, but only in the CD8^HIGH^ *PDCD1* (PD-1)^HIGH^ group. In fact, where IHC stratification of PD-L1 expression has failed to predict immunotherapeutic responses, KLRC1 stratification in IMvigor210 showed protective effects were restricted to the PD-L1 IC high group.

Combined NK and T cell strategies are actively being explored in existing clinical trials, such as the COAST trial (NCT03822351) of combination anti-PD-L1 anti-NKG2A in stage III unresectable nonsmall cell lung cancer. Interim results show that combination therapy provides substantial gains in achieving a 10-month progression-free survival rate of 72.7%, compared to only 39.2% in the durvalumab-only arm (57). Our data support the combination immunotherapeutic approach, indicating that BCG-induced adaptive immune resistance raises PD-L1 levels alongside HLA-E on tumor cells. While this analysis does not exclude the presence of alternative checkpoints, such as TIGIT, TIM3, or LAG3, it lends evidence to the hypothesis that combination NK and T cell immunotherapeutic approaches may hold the key to improved outcomes for NMIBC patients.

## Conclusions

Our data suggest that inflammation was ubiquitous in all patients after BCG immunotherapy, regardless of their recurrence status, and that IFN-γ directly contributed elevation in HLA-E and PD-L1 expression by recurring tumors in the TME. We found that NKG2A expression stratified NK and T cells by cytolytic capacity, although NKG2A-expressing cells additionally presented secondary resistance checkpoint receptors. *In situ* spatial analyses revealed that HLA-E^HIGH^ tumors are activated to recruit NK and T cells via chemokine production, potentially sparing HLA-E^LOW^ tumors that would otherwise be susceptible to lysis. Finally, blood-derived NK cells retained anti-tumor functions at the time of tumor recurrence. These data support the potential of combined checkpoint blockade, specifically NKG2A and PD-L1, as a potential bladder-sparing mechanism for overcoming BCG-induced immune exhaustion.

## Methods

### Patients and samples

Patients at Mount Sinai Hospital (MSH) were enrolled in the study following Institutional Review Board (IRB) approval (protocol 10-1180). 10-1180 covers the use of patient tissues in a biorepository and allows for prospective collection of blood, urine, and tissue samples from enrolled patients. Formalin-fixed paraffin-embedded (FFPE) blocks from BCG patients were obtained retrospectively from the biorepository and prospectively for patients receiving treatment. For prospective patients, samples were collected on the day of surgery and throughout BCG immunotherapy. Due to IRB limits on the collection, blood and urine samples were taken at the first dose, third dose, and sixth dose of the induction cycle; at every follow-up cystoscopy; and at the third maintenance cycle dose. Tumor samples were taken at every possible timepoint. BCG naïve was defined as any patient who had yet to receive BCG, regardless of past treatment with other chemotherapies. BCG-unresponsive was defined as any patient with recurrent tumors following at least five of six induction doses of BCG.

### Sample processing

Blood and urine samples from bladder cancer patients were processed to collect PBMCs, serum, and cell-free urine. Blood was spun down (4C, 2000rpm, 10 min) to isolate serum, and PBMCs were collected using Ficoll-Paque isolation. PBMCs were frozen down in a Mr. Frosty at −80 C for 24 hours and stored at −160 C in 10% DMSO and 90% fetal bovine serum; PBMCs and cell-free urine were stored at −80 C.

Tumor tissues obtained from transurethral resections of bladder tumor (TURBT) were placed into RPMI medium immediately after removal and transferred to the laboratory for additional processing. Bladder and lymph nodes obtained from radical cystectomies were sent from the operating room directly to the pathology suite after completion of the lymph node dissection. The bladder was bivalved; samples of visible tumor were extracted and placed in RPMI medium, and the tumor was transferred to the laboratory for additional processing.

Fresh tumor samples underwent a variety of different processing techniques based on the planned experiment. Tumor used for spatial sequencing was placed in a 10 mm x 10 mm cryomold with optimal cutting temperature (OCT) media and frozen down on a thermal block immersed in liquid nitrogen. Tumor tissues were digested using tumor dissociation enzymes (Miltenyi, 130-095-929) and a GentleMACS machine (program 37C_h_TDK_3) at 37C. Mechanically and enzymatically separated tissues were filtered through a 70μM cell strainer and assessed on Countess II (ThermoFisher) for viability and cell numbers.

### Targeted RNA sequencing

FFPE sections from 38 retrospective NMIBC cases were obtained from the institutional biorepository and used for targeted RNA sequencing. RNA was extracted from five and ten μm sections. HTG EdgeSeq lysis buffer was added to lyse and permeabilize the samples. Nuclease protection probes (NPPs) were added to the lysed samples and hybridized to the target mRNA. A nuclease was added to digest non-hybridized mRNA and excess NPPs. The nuclease digestion reaction was finalized with a termination solution followed by heat-mediated denaturation of the enzyme.

Each sample was used as a template for PCR reactions with specially designed primers. Each primer contains a unique barcode that is used for sample identification and multiplexing. Samples were analyzed simultaneously on an Illumina sequencing platform to prepare the library. All samples and controls were quantified in triplicates. No template control (NTC) reactions were made for each master mix used during the qPCR process to test the absence of a probe or qPCR contamination. Molecular-grade water was used in place of a test sample in the NTC reactions using the same volume as the template.

Sufficient concentration of sample for library pooling, appropriate dilution for the library pool, and volume of denaturation reagents to add to the library were determined by HTG library calculator. 2N NaOH and heat (98C, 4 minutes) were used for library denaturation. The denatured library was loaded into the well of the NextSeq sequencing cartridge. Sequencing was performed using an Illumina NextSeq sequencer.

The sequencing data on mRNA expression of target genes were imported into HTG EdgeSeq parser software. HTG biostatistics department performed quality control analyses and normalized the data. Data were returned from the sequencer as demultiplexed FASTQ files with four files per assay well. The HTG EdgeSeq parser software aligned the FASTQ files to the probe list and collated the data.

### Protein concentration measurement

Cell-free urine supernatant and serum samples were randomized in a 96-well plate. Diluted samples and positive and negative controls were incubated overnight with an incubation mix (incubation solution, incubation stabilizer, A-probes, and B-probes) at 4°C. Samples were then incubated with an extension mix (High purity water, PEA solution, PEA enzyme, PCR polymerase) for 5 min and placed on a thermal cycler. Following the thermal cycler, samples were incubated with a detection mix (detection solution, High purity water, detection enzyme, PCR polymerase) and transferred to a chip. Primers were loaded onto the chip, and the chip was profiled using the Fluidigm IFC controller HX with the Fluidigm Biomark Reader. Data were normalized using extension and interplate controls and a correction factor. The resulting data were reported in normalized protein expression (NPX) units on a log2 scale.

### IFN-γ co-culture

Cell lines and CD45-isolated primary tumor cells were incubated in media optimized for high viability for 72 hours (RPMI-1640 supplemented with 20% fetal bovine serum). Tumor cells were expanded until they were confluent in 2 T175 flasks. Following expansion, cells were culture in 100 ng / mL of IFN-γ for a total of 24 hours in a 24 well plate. Following co-culture, cell lines were trypsinized (immortalized), and primary tumors were gently removed from the solid phase by a cell scraper. Samples with and without IFN-γ stimulation were measured for HLA-E and PD-L1 protein expression levels.

HLA-E and PD-L1 levels were assessed via FlowCytometry. Cells were stained in 4C FACS buffer (phosphate-buffered saline (PBS) with 2% heat-inactivated FBS and EDTA 2 mM) for 30 minutes. Subsequently, cells were washed in PBS, incubated for 20 minutes in a viability dye, washed again with PBS, and resuspended in 2% paraformaldehyde. The experiment was performed in triplicate, with three readouts per cell line per experimental condition. FlowCytometry acquisition was performed using an LRS-Fortessa (BD Biosciences), and data were analyzed using the CytoBank software. When staining for HLA-E, cells were first stained 20 minutes with HLA-E prior to staining with additional PD-L1. In CytoBank, several gates were applied to generate the final dataset. A live/dead gate was applied, followed by a gate to remove doublets and isolate singlets. Lastly, the data was arcsinh transformed prior to analysis.

Once the final dataset had been generated, statistical significance between unstimulated and stimulated cell lines was assessed. An independent t-test was run with a significance cutoff of p < 0.05.

### Targeted RNA sequencing analyses

Prior to any analyses, clinical data on all patients was gathered. The following items were collected for all samples processed by HTG Molecular: patient date of birth; gender; age at BCG induction; BCG status (unresponsive vs. naïve vs. exposed); date of sample collection; stage at collection; grade at collection; recurrence date; time to recurrence; time to progression; BCG start date; BCG last exposure; BCG induction cycle last dose; prior chemotherapy; cystectomy; date of cystectomy; stage at cystectomy.

Following clinical data collection, a gene set enrichment analysis (GSEA) was performed on the targeted RNA sequencing data. Specifically, we used paired patient samples before and after BCG exposure in the BCG recurrent patient population only. We used custom gene sets, as well as all Hallmark gene sets from the Broad Institute’s MSigDB, as inputs for the enrichment analysis (24). Statistical significance was set at p < 0.05, and gene sets found to be significant are listed in **Supplementary Table S1**. All gene sets found to be statistically significant were evaluated for leading-edge genes, defined as the genes that contribute most to the enrichment score and associated p-value.

The leading-edge genes from statistically significant gene sets in the GSEA were collated and used to assess for group differences between the paired HTG patient samples. We performed these analyses specifically on the BCG-recurrent cohort. Prior to any analyses, a Shapiro-Wilk test, chosen for suitability in small sample sizes, was used to assess for normality (58). All samples with p < 0.05 were considered not normally distributed, and a Kruskal-Wallis test was performed to assess for group differences (59). All other samples were assessed using an independent T-test. Genes with statistically significant differences between the BCG naïve and the BCG-unresponsive populations were then visualized on radar plots.

### Plasma and cell-free urine supernatant protein concentration analysis

We used the OLINK Proteomics^®^- inflammation panel to validate the signature seen in the Targeted Bulk RNA sequencing data. Protein levels were visualized at BCG naïve, third dose, sixth dose, and first-cystoscopy timepoint, comparing NPX values for all patients between the BCG naïve timepoint and the sixth dose timepoint. In order to determine the suitable statistical test, a Shapiro-Wilk’s test was used to assess for normality, and a Kruskal-Wallis test was used in every instance in which one or both samples were not normally distributed. An independent T-test was used in the event both samples were normally distributed. All statistically significant p values were then used to assess adjusted p values via the Benjamini-Hochberg correction, with an alpha of 0.5. All statistically significant genes between the BCG naïve and sixth induction dose time points are shown.

### Single-cell RNA sequencing

#### Data preprocessing

Single-cell RNA sequencing (scRNA-seq) analysis was performed using Seurat v4.0.1 in RStudio v1.1.456 with R v4.0.3 (60). Genes expressed in fewer than three cells were excluded from later analyses. Cells with < 200 or > 2500 unique genes, as well as cells containing >15% mitochondrial gene transcripts, were discarded. Subsequent data normalization was performed using the NormalizeData function and consisted of dividing feature counts for each cell by total counts for that cell, scaling by a factor of 10,000, and natural log transformation. Two thousand variable genes in each sample were identified using the FindVariableFeatures function. We then integrated 24,166 cells containing 26,520 genes from five bladder tumor samples via the IntegrateData function and performed scaling and principal component analysis (PCA) on the integrated data structure using the functions ScaleData and RunPCA, respectively. Using the first 50 principal components (PCs), graph-based clustering was performed using Seurat FindNeighbors and FindClusters functions (resolution = 0.9), and UMAP dimensionality reduction was performed to reveal 28 cell clusters. Cluster-specific marker genes were identified using the FindAllMarkers function, and marker genes with a natural-log fold change (FC) > 0.25 and expressed in >= 25% of cells were used to annotate cell cluster identities based on known cell type markers (61–63): CD8 T cells (*CD8A*, *CD3E*), CD8-T cells (*CD3E*), tumors (*MUC1, KRT7/8/13/17/18*), B cells (*MS4A1, CD79A*), monocytes/macrophages (*CD14, LYZ, FCGR3A, MS4A7*), T regulatory cells (*CD3E, FOXP3*), granulocytes (*C1QA/B/C, ITGAM, FCGR3A*), NK (*NKG7*, *GNLY, KLRD1, KLRF1*), fibroblasts (*COL1A1, SPARC*), epithelium (*EMP1*), DC (*FCER1A, CST3, ITGAX*), plasma cells (*MZB1, JCHAIN, IGHG3*), and CD8 TRM cells (*CD3E*, *CD8A, KLRD1*). Clusters 22, 24, 25, and 26 containing 224, 142, 141, and 101 cells, respectively, were ultimately excluded from further analyses due to unclear cell identities.

#### NicheNet Analysis

We used the NicheNet R package to infer potential ligand-receptor interactions between NK and T cells and tumor cells, as well as potential regulatory effects in tumor gene expression as a result of these interactions (31). We added known ligand-receptor interactions between ligands IgG1 and IgG3 and the receptor CD16 (weight = 1) to NicheNet’s default ligand-receptor interactions database, and also integrated the Harmonizome Pathway Commons Protein-Protein Interactions database: (https://maayanlab.cloud/Harmonizome/dataset/Pathway+Commons+Protein-Protein+Interactions) with NicheNet’s default signaling network (weight = 1) (64,65). We used the modified ligand-receptor interaction and signaling networks for subsequent analyses. We defined the NK and T cell populations as senders and tumor cells as receivers and derived a list of 92 receptors expressed in the receiver population by intersecting the list of receptors from the ligand-receptor interaction network with genes expressed in at least 10% of tumor cells in the RNA assay. Fifty potential ligands expressed in greater than 10% of the sender NK and T cells were then derived based on their interaction with the list of 92 receptors. We then calculated activity scores for each ligand based on its potential to affect the expression of Hallmark interferon gamma response genes (24,66). Statistical analyses including Wilcoxon rank-sum tests and Pearson and Spearman correlations were performed using R v4.0.3.

#### Differential gene expression analysis

Differentially expressed genes (DEGs) between KLRC1^+^ versus KLRC1^-^ cells were found using the FindMarkers function from the Seurat R package, where cells with KLRC1 expression greater than 0 in the RNA assay were labeled as KLRC1^+^.

#### 10X Visium spatial sequencing

Tumor samples collected both prospectively and retrospectively were used to generate a potential cohort for spatial sequencing. A 10 μm slice of tumor was hemotoxin and eosin-stained (H&E) and reviewed with a board-certified pathologist to verify tissue integrity, the presence of tumor nests, and immune infiltrates. Three - five 10 μm slices from each tumor were then used for RNA extraction and quantification. A Qiagen RNEasy Mini kit was used for RNA extraction; a ThermoFisher NanoDrop was used to assess extracted RNA concentration; and an Agilent TapeStation was used to identify RNA integrity via the RNA integrity number equivalent (RINe) score. A RINe cutoff of > seven was used as a threshold for good quality.

Of an initial cohort of 55 samples, eight were identified as having good-quality tissue integrity, presence of tumor / immune populations, and high RNA quality. One representative sample from the cohort was used for tissue optimization, which generates the optimal experimental parameters for the spatial sequencing cohort. Tissue optimization consists of 1) tissue permeabilization, 2) fluorescent cDNA synthesis, 3) tissue removal, and 4) slide imaging. Tissue slices are cut onto the capture areas which contain poly(dT) primers for the capture and production of cDNA. After the addition of the master mix containing fluorescently labeled nucleotides, fluorescent cDNA is attached, and can be visualized following enzymatic removal of the tissue. After 7 variable timepoints from 5 minutes to 60 minutes, the optimal duration of permeabilization is determined by the timepoint with the highest fluorescence signal and the lowest signal diffusion.

Using the parameters generated by the tissue optimization step, a cohort of 8 samples was used for the full experimental protocol. Tissues were stained and imaged; cDNA was synthesized; a second cDNA strand was synthesized and denatured; cDNA was generated and the first quality control was performed. Finally, the Visium Spatial gene expression library was constructed, and the second QC was performed. Following library construction, the samples were sequenced and aligned using Illumina platform.

#### Spatial neighbors, differential gene expression analyses

Spatial sequencing analyses were performed in Python 3.8.8 using Scanpy, Squidpy, AnnData, and the Tangram packages. Data storage and computational job submission were run on Mount Sinai’s high-performance computing cluster, Minerva. Each spatial object was normalized, log-transformed, and filtered. Visium dots with fewer than 1,000 and more than 35,000 transcribed genes were filtered out. Next, the resulting spatial objects were concatenated into a single object, and genes were filtered out if they appeared in less than 1% of the total number of Visium spots. Labeled scRNA-seq data was then used to project cell-type annotations onto the spatial data in a multi-label multi-class classification algorithm (Tangram). Tangram assigns probabilities that a specific cell type resides in a given Visium spot. Visium spots have a diameter of 55 μm, and therefore multiple cells can reside in a single spot, or “nano-environment.” A spot was designated as having a given cell type if the mapped single-cell probability in that spot was > 50%. Lastly, clinical annotations associated with the samples were added to the spatial data to create a complete cohort.

Following projection of cell-type annotations, Tregs were labeled according to canonical lineage markers. Labeled CD4+ T cells were assessed for positivity of the following markers derived from the literature: *CCR8, CCL1, IL-10, TGFB1, ENTPD1, CTLA-4, IL-2RA, LAG3, LAYN*, and *CCR8*. Of these, *TGFB1, ENTPD1*, and *LAYN* were present in our dataset. All CD4+ T cells with expression of these 3 genes were considered Tregs.

Spatial relationships between tumor cells, immune infiltrates, and relevant genes were then compared visually and numerically. *HLA-E* expression was binarized for all Visium dots into those expressing any level of *HLA-E* and those expressing no *HLA-E*. These binarized data were used to create topographical density plots of HLA-E expression (Fig. 3). *HLA-E* quartiles were computed, and high *HLA-E* expressing cells were designated as those > 75^th^ percentile of expression. Lastly, *IFNGR1* and *IFNGR2*, and *CXCL9/10/11* gene scores were created by scoring expression against 150 randomly selected genes to create a single numerical score.

A spatial neighbors’ analysis was computed using a radius of 2 Visium dots. Each Visium dot was therefore defined as a function of its surrounding labeled Visium dots, out to 2 rings adjacent. Visium dots containing tumor cells were isolated, and each tumor-containing dot was characterized as a function of its surrounding tumor-infiltrating T cells. A differential gene expression was run, comparing gene expression of the Visium dots with no surrounding infiltrating tumor cells to cohorts of 1 and greater surrounding T cells. An independent T test was used, and the Benjamini-Hochberg correction was applied to generate adjusted P values.

Heatmaps were generated to visualize gene expression as a function of the neighboring nano-environment. Heatmaps were generated using row-scaled expression (from 0 – 1) for heatmaps with greater than two columns, and Z-scored expression (for heatmaps with two columns). For both CD4+ infiltrates and NK/CD8+ infiltrates, we generated cohorts that were HLA-E^HIGH^ and infiltrate^HIGH^ cases. For both conditions, HLA-E was considered high if expression surpassed the 75^th^ percentile. For CD4+ T cells, infiltrates were high of there were seven or more neighboring CD4+ cells; infiltrates were low if there were four or fewer neighboring CD4+ cells. For cytolytic cells, infiltrates were high if there were one or more neighboring NK or CD8+ T cell, and low otherwise.

Lastly, a distance experiment was performed to characterize each tumor-identified Visium spot as a function of the closest cytolytic cell or Treg. The distance between all tumor Visium dots and all cytolytic or Tregs on a slice-by-slice basis was computed using scipy’s cdist function, which calculates the Euclidian distance between two points. Then, having a matrix of tumor cells in the rows, cytolytic or Treg cells in the columns, and the distances between them as values, the minimum of each row was taken. This data was stored as metadata in the spatial objects. Histograms colored by BCG exposure status were created. Scatterplots with *CXCL9/10/11* on the y-axis and distance to the most proximal cell of interest on the x-axis were generated.

#### Mass cytometry antibody preparation and staining

PBMCs from BCG-treated bladder cancer patients at the time of tumor recurrence were isolated using Ficoll-Paque and resuspended in cell medium (RPMI-1640 medium supplemented with 10% heat-inactivated FBS, 1% Penicillin, 1% Streptomycin and 1% L-glutamine). PBMCs were conjugated to antibodies purchased from Fluidigm using the Maxpar X8 and MCP9 labeling kits. Platinum barcodes were prepared as previously described. All antibodies were titrated prior to conjugation.

Prior to mass cytometry staining, all cells were incubated for 20 minutes at 37C in RPMI cell medium (described above) and IdU (Fluidigm, t#201127), Rh103 (Fluidigm, #201103A) and anti-IgG4 if cells were to be co-cultured with K562 and rituximab. Following the incubation, cells were centrifuged, washed with PBS and 0.2% bovine serum albumin (BSA), and incubated for 3 minutes on ice with an Fc-blocking reagent. Samples were washed again with PBS and 0.2% BSA; barcoded samples were pooled together and washed; finally, samples were stained with extracellular antibodies for 30 minutes on ice and in PBS and 0.2% BSA.

When staining four samples, cells were single-barcoded using 194Pt, 195Pt, 196Pt or 198Pt. When staining five or six samples, cells were stained with combination of barcodes. Cells were again washed using PBS 0.2% BSA; barcoded samples were pooled together; samples were washed again, and stained with extracellular antibodies for 30 minutes on ice in PBS 0.2% BSA. Cells were co-cultured in the presence of either K562 and rituximab, Raji cells, or nothing (Med condition), for a total of 6 hours.

Samples were washed with PBS 0.2% BSA and resuspended in Fixation/Perm buffer (Invitrogen, #00-5523-00) for 30 minutes on ice. Cells were centrifuged and washed with Maxpar Barcode Perm Buffer (Fluidigm, #201057) and barcoded using the Cell-ID 20-Plex Pd Barcoding kit (Fluidigm, #201060). Barcoded samples were washed with permeabilization buffer (Invitrogen, #00-5523-00) and pooled. Intracellular staining was then performed in permeabilization buffer with Heparin at a concentration of 100U/mL for 30 minutes on ice. Stained cells were washed with permeabilization buffer and resuspended in PBS with PFA 2.4%, saponin 0.08%, Osmium tetroxide 0.075nM and Ir 0.125uM (Fluidigm, #201192A). Finally, samples were washed and resuspended in PBS 0.2% BSA and data were acquired within four days, or frozen in FBS/DMSO 90/10. The antibody panel in Supplemental Table S2 was used to stain bladder cancer patient samples.

### Mass cytometry sample acquisition and processing

Prior to acquisition, samples were washed with cell staining buffer and acquisition solution (Fluidigm). Following washing, samples were resuspended in acquisition solution (1 million cells / 1 mL) containing a 1:20 dilution of EQ normalization beads. Data were then acquired using the Fluidigm Helios mass cytometer with a wide bore injector configuration at an acquisition speed of < 400 cells per second. The output files were normalized and concatenated using Fluidigm’s CyTOF software, and outputted as FCS files.

The Mount Sinai Human Immune Monitoring Core’s (HIMC) pipeline for processing and cleaning was used to clean the resulting FCS files. Aberrant acquisition time-windows and low DNA intensity events were stripped out by the sample preprocessing pipeline. Samples were then demultiplexed via the cosine similarity of the Palladium barcoding channel on a cell-by-cell basis to every possible barcode used in a batch. Once the cell-barcode labeling has been established, the signal-to-noise (SNR) ratio was calculated by taking the difference between the highest and second highest similarity scores. Cells with low SNR ratios were flagged as multiplets and removed. Finally, acquisition multiplets are removed based on the Gaussian parameters residual and offset acquired by the Helios mass cytometer.

### Data processing and analysis

Data were uploaded onto Cytobank and processed for downstream analyses. Several gates were applied: a live dead gate and a doublets gate in sequential order. All data were arc-sinh transformed, and no batch corrections were performed given all samples were run in a single batch. NK cells were identified via manual gating assignment. Files were downloaded onto Minerva, concatenated into a single object, and clinical data were assigned to each sample. A phenograph analysis was performed to cluster the cellular data. A MELD analysis was run stratifying samples at the tumor recurrent timepoint between the ex-vivo state and the K562 or Raji cell states. Highly polarized clusters (<25% average predicted likelihood of being in the ex-vivo state; > 75% average predicted likelihood of being in the Raji or K562 state) were isolated, and a differential protein expression was run. Wilcoxon tests were run for statistical significance and the Benjamini-Hochburg correction was applied.

## Data Availability Statement

data were generated by the authors and will be uploaded to the Gene Expression Omnibus upon publication of the article.

