## Supplemental tables and figures for "Elevated HLA-E and NKG2A as a Consequence of Chronic Immune Activation Defines Resistance to *M. bovis* BCG Immunotherapy in Non-Muscle-Invasive Bladder Cancer" (from another version of this preprint: version 2, 2022-03-31)

| Gene Set Names | Defining Genes |
|---|---|
| Interferon Gamma Signature | *CCL5, CD3D, CXCL10, IL2RB, LAG3, IDO1, IL2RG, STAT1, PRF1, PDCD1, GZMA, GZMB, CD8A, CXCL9, TBX21, CD27, CD4, CD74, IFNG, CCR5, CXCL11, CXCL13, CXCR6, PTPRC* |
| B Cell Markers | *CD79A, BLNK, CD79B, CCR9, CD19, PNOC, BLK, CD22, HLA-DOB, TCL1A, CR2, SPIB, CD72, TNFRSF17, HLA-DQA1, PAX5, MS4A1, MEF2C, FCRL2* |
| Cytotoxic CD8 T-Cells | *GZMK, FLT3LG, KLRK1, KLRD1, PRF1, GZMA, CD3G, CD8B, SIGIRR, GZMB, KLRB1, CD8A, GZMH, PF4, TBX21, CD3E, NKG7, CD27, IFNG, GNLY, EOMES, GZMM, KLRF1, RUNX3* |
| Cytotoxic Receptors and Function | *ZNF683, SLAMF6, KLRK1, KLRD1, SLAMF7, NCR1, SH2D1B, GZMA, TYROBP, SH2D1A, GZMB, KLRB1, KIR3DL3, CLEC4C, CLEC4D, GZMH, NKG7, NCR3, GNLY, GZMM, KLRF1* |
| T-Cell Checkpoint Modulation | *HLA-DQA2, CD274, TNFRSF14, CD86, CARD11, CD40, AKT1, CD3D, TNFRSF9, KLRK1, LAG3, CSK, TNFRSF13C, CD276, CD80, PDCD1, CD40LG, CD3G, PDCD1LG2, PTPN6, HLA-DQB2, ADORA2A, HLA-DPB1, TNFSF4, TIGIT, HLA-DRB1, CCL19, HAVCR2, CD3E, CD27, CD5, CCL21, CD4, HLA-DQA1, VAV1, CD70, CD24, GRB2, SRC, HLA-DRA, CD28, TNFSF13B, HLA-DPA1, TNFRSF4, CTLA4, ICOS, LYN, BTLA* |
| Tumor Inflammation | *PRF1, GZMA, GZMK, CCL5, CXCL10, IFNG, CXCL9, STAT1* |
| Th1 Polarization and Response | *NFKB1, RELA, IL12B, TBX21, IL2, IFNGR1, IL12RB1, RBPJ, JAK2, TYK2, JAK1, DLL1, IFNG, STAT1, STAT4, IL12A* |
| Lymphocyte Trafficking | *CD274, FAM129C, CDH2, CXCR4, CD68, PDCD1, PDCD1LG2, PLCG2, CD40LG, HLA-DOB, MAPK11, ITGAL, CD52, HLA-DPB1, HLA-DRB1, PVR, CD4, ITGA5, HLA-DQA1, VAV1, NCAM1, HLA-C, LILRB3, CTLA4, CD83, CD86, LILRB1, HLA-DOA, LILRA3, CXCL12, ROCK2, LILRA5, ITGAX, CD8A, PLCG1, TIGIT, HLA-DMB, ITGA4, MMP9, CD28, CYBB, SELE, ICOS, CLEC4E, ICAM1, HLA-DQA2, SDC1, CEACAM3, CD96, CD80, CD22, CD84, SELL, CD209, PDPN, SIGLEC5, HLA-DRA, SIGLEC1, ITGB2, S1PR4, CD7, CD177, ITGAM, HLA-A, ITK, CD40, ITGB7, CD34, MSN, HLA-F, CD276, PTK2B, RASSF5, CD6, S1PR1, MAPK14, VCAM1, HLA-E, TREM1, HLA-DMA, HLA-DPA1* |
| General T Cell Markers | *LCK, CD247, CD2, CD3E, CD3G, CD5, SH2D1A, TNFRSF25, TRAT1, CD96, CD6, LAMP1, CD3D, IL2RB, CD14, ZAP70* |
| Receptor | *PDCD1, HAVCR2, TNFRSF14, CD27, CD40, TLR4, KIR2DL1, ICOS, BTLA, TNFRSF9, ADORA2A, LAG3, TNFRSF4, CTLA4, TIGIT, IL2RA, TNFRSF18* |
| Stress Toxicity | *RAD17, DDIT3, TLR4, ATR, ADM, BID, CHEK2, HSP90B1, IL1B, RIPK1, CCL2, IL6, HSPA5, CD40LG, CRP, GADD45A, VEGFA, DDB2, HSP90AA1, CASP1, TNFRSF10A, PARP1, PVR, CDKN1A, TP53, TNFRSF10B, NBN, EDN1, FAS, SERPINE1, RAD51, NFAT5, ATG7, CA9, IFNG, ATG5, ATG12, XPC, CALR, GRB2, RAD9A, HMOX1, MMP9, HUS1, EPO, IL1A, CHEK1, ATM, TNF* |
| DNA Repair (Hallmark) | *TYMS, POLR1D, GTF2H1, ITPA, TAF6, TK2, SDCBP, GPX4, SMAD5, SAC3D1, UMPS, NUDT9, NME3, RPA3, NFX1, DGUOK, GTF2H3, DGCR8, DCTN4, ERCC2, TAF9, POLR2H, NME4, XPC, BOLA2_2B, DAD1, VPS37D, TMED2, NCBP2, RAE1, NUDT21, CCNO, GTF3C5, ERCC4, SUPT5H, ARL6IP1, EIF1B, ZNF707, AK3, VPS28, RAD52, TAF10, ERCC3, FEN1, SUPT4H1, PCNA, POLR2A, POLH, BCAP31, POLB, NT5C, HCLS1, SRSF6, POLR2J, REV3L, USP11, VPS37B, POLR2K, GTF2H5, ADCY6, HPRT1, TP53, PRIM1, RALA, BCAM, POLD1, CLP1, SURF1, POLD4, GMPR2, MPG, PNP, DDB1, POLR2D, RBX1, SF3A3, RFC4, ZWINT, IMPDH2, POLR2C, ADRM1, PDE6G, POLR2G, POLR1C, POLR2F, LIG1, DUT, CMPK2, SEC61A1, RFC3, GUK1, PDE4B, RAD51, GTF2F1, POLA2, STX3, TARBP2, CDA, APRT, ELL, RFC5, POM121, RPA2, POLR3C, NME1, POLR2E, AK1, NPR2, ERCC1, GSDME, TAF12, POLR2I, POLA1, CETN2, POLL, ZNRD1, POLD3, RNMT, TSG101, ELOA, DDB2, UPF3B, SNAPC5, CANT1, ADA, POLR3GL, TAF13, COX17, ERCC8, AAAS, BRF2, POLE4, SSRP1, TAF1C, RFC2, SNAPC4, EDF1, ERCC5, CSTF3, GTF2B, GTF2A2, NELFCD, MRPL40, RRM2B* |
| P53 Pathway (Hallmark) | *LRMP, IRAK1, DDIT3, AK1, BAK1, ACVR1B, RGS16, ITGB4, IFI30, HBEGF, SDC1, APAF1, MXD1, SOCS1, TXNIP, VWA5A, TRAFD1, BTG2, RAD51C, CEBPA, SPHK1, DGKA, HRAS, BAX, DDB2, GADD45A, CD82, TGFB1, RRAD, ELP1, FBXW7, CYFIP2, TCHH, PTPRE, ADA, PCNA, CDH13, CD81, CASP1, POLH, AEN, SLC3A2, SEC61A1, CDKN1A, LIF, TP53, PHLDA3, FAS, NINJ1, DEF6, FOS, TSPYL2, XPC, PROCR, HDAC3, RAD9A, HMOX1, ERCC5, CCND2, POM121, FOXO3, IL1A, CCND3, TNFSF9, TRAF4, TAP1, APP, HIST1H1C, PPP1R15A, NOTCH1, BTG1* |
| Oxidative Phosphorylation (Hallmark) | *VDAC1, ALAS1, HSD17B10, SDHA, ALDH6A1, ATP5F1, PHB2, BAX, TCIRG1, GPX4, CASP7, NDUFA5, POLR2F, COX17, HTRA2, SURF1, OAT, SLC25A3, DLAT, GPI* |
| Peroxisome (Hallmark) | *HSD11B2, ABCD2, ABCB1, ERCC1, ERCC3, HRAS, CDK7, CADM1, FDPS, BCL10, ACSL1, MSH2* |
| Hypoxia | *NFKB1, VDAC1, ADORA2B, ATR, NAMPT, ADM, TXNIP, MAP3K1, F3, VEGFA, EGR1, RBPJ, MIF, IGFBP3, NCOA1, HIF1A, PLAU, TP53, EDN1, SERPINE1, FOS, CA9, PER1, NOS3, MMP9, HMOX1, LGALS3, MET, EPO, APEX1, BLM, GPI, PIM1* |
| TGF Beta Signaling | *CBL, CREBBP, LTBP1, ID1, TGFB1, SMAD5, MYC, PARP1, TGFBR3, EP300, IFNG, TGFBR1, SMAD2, TGFBR2, THBS1, SMAD3, TGFB3, MAPK3, TNF* |

**Supplementary Table S1:** Genes defining gene-set enrichment pathways

| Cell Type | Cell # | Cluster(s) |
|---|---|---|
| NK Cells | 1035 | 14, 21 |
| T Cells | 2529 | 5, 9 |
| CD8+ T Cells | 2136 | 1 |
| CD8+ TRM Cells | 223 | 23 |
| T Reg Cells | 973 | 10 |
| B Cells | 1269 | 8 |
| Plasma Cells | 360 | 20 |
| Monocyte/Macrophage | 1333 | 6 |
| DCs | 1827 | 2 |
| Granulocytes | 1759 | 7, 19 |
| Fibroblasts | 916 | 11 |
| Epithelium | 487 | 16 |
| Tumor Cells | 8711 | 0, 3, 4, 12, 13, 15, 17, 18, 27 |
| 22 | 224 | 22 |
| 24 | 142 | 24 |
| 25 | 141 | 25 |
| 26 | 101 | 26 |

**Supplementary Table S2:** Cell numbers and identities associated with the scRNA sequencing clustering analysis

| From | To | Weight |
| --- | --- | --- |
| IFNG | HLA-E | 0.201 |
| MYC | HLA-E | 1.5 |
| NFKB1 | HLA-E | 1.14 |
| STAT1 | HLA-E | 1.98 |

**Supplementary Table S3:** Niche-Net predicted weights between IFNg and downstream signaling mediators

| Metal | Conjugated Antibody |
|---|---|
| 89 Y | CD45 |
| 111 Cd | HLA-DR |
| 112 Cd | Granzyme A |
| 113 Cd | CD38 |
| 114 Cd | CD11B |
| 115 In | IFN-Y |
| 116 Cd | CD57 |
| 141 Pr | cKIT |
| 142 Nd | KLRG1 |
| 143 Nd | Granzyme K |
| 144 Nd | CD69 |
| 145 Nd | NKG2D |
| 146 Nd | DNAM1 |
| 147 Sm | NKp80 |
| 148 Nd | KIR3DL1 |
| 149 Sm | KIR3DL1 |
| 150 Nd | IL-2 |
| 151 Eu | CD107a |
| 152 Sm | TNF-a |
| 153 Eu | CD14 |
| 154 Sm | MIP1b |
| 155 Gd | NKp46 |
| 156 Gd | Tim-3 |
| 158 Gd | KIR2DL1 |
| 159 Tb | CD56 |
| 160 Gd | NKG2A |
| 161 Dy | NKP44 |
| 162 Dy | CD27 |
| 163 Dy | EOMES |
| 164 Dy | NKG2C |
| 165 Ho | KIR2DL3 |
| 166 Er | KIR2DL1_S1 |
| 167 Er | KIR2DL2_L3 |
| 168 Er | TRAIL |
| 169 Tm | Tbet |
| 170 Er | CD3 |
| 171 Yb | CD127 |
| 172 Yb | Perforin |
| 173 Yb | Granzyme B |
| 174 Yb | TIGIT |
| 175 Lu | XCL1 |
| 176 Yb | CD200R |
| 209 Bi | CD16 |

**Supplementary Table S3:** Metals and antibodies used for CYTOF

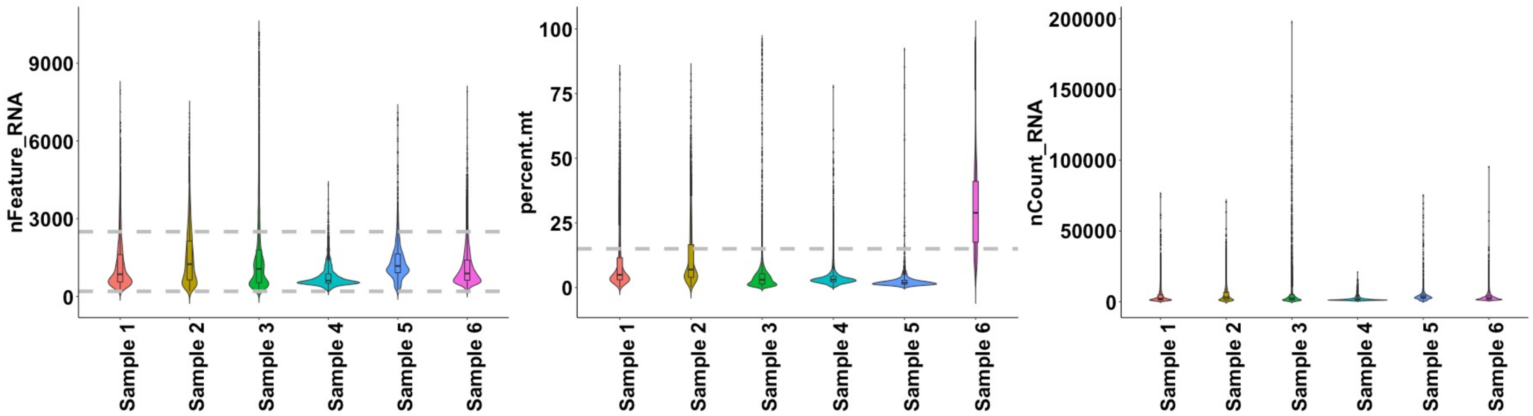


**Supplementary Figure S1:** QC plots for scRNA-Seq samples used. Violin plots demonstrating number of unique features (left), percent mitochondrial genes (middle), and total feature count (right) for each sample used in subsequent scRNA-Seq analyses.

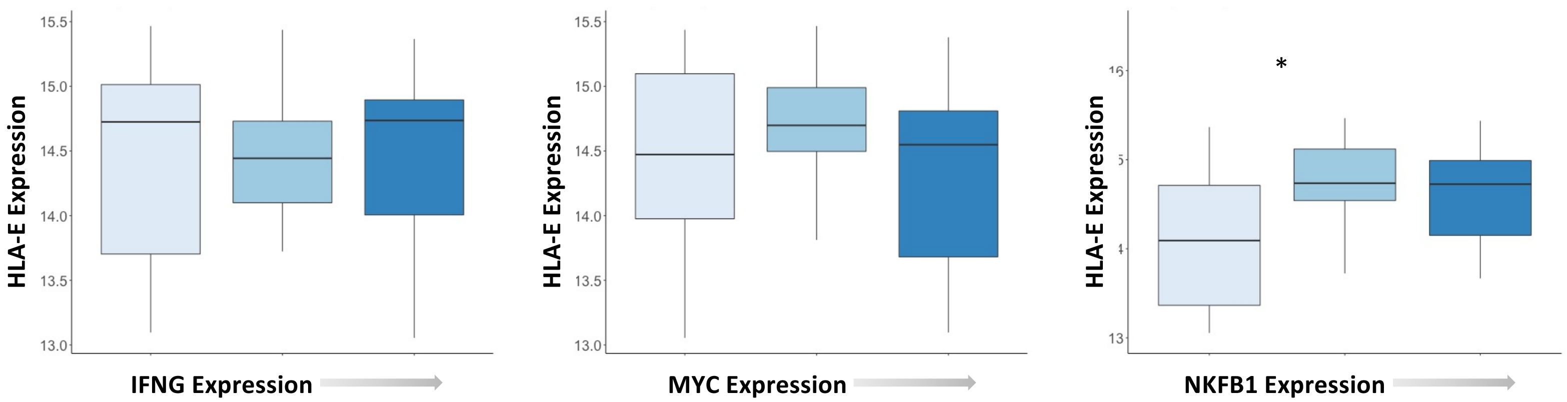


Expression of HLA-E in HTG bulk RNA-Seq urothelial cancer cohort with respect to IFNG (left), MYC (middle), and NFKB1 (right). levels. (*p = 0.044)

**Supplementary Figure S2:** *HLA-E* expression in HTG bulk RNA-Seq bladder tumor cohort relative to *IFNG*, *MYC*, and *NFKB1* levels.

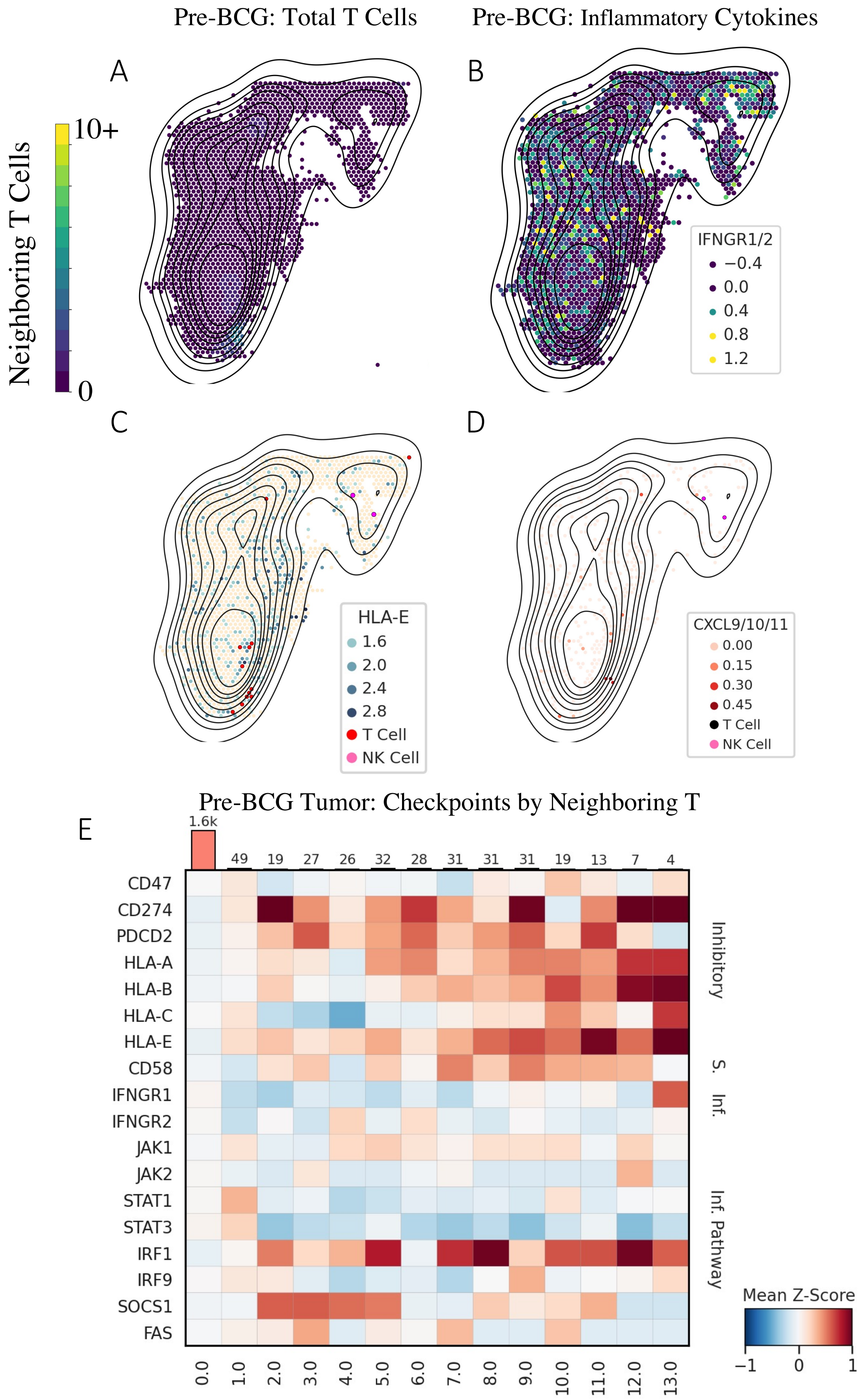


**Supplementary Figure S3:** Visualization of spatial sequencing in a representative post-BCG sample. **A:** a representative pre-BCG slice showing neighboring T cell infiltrates (both CD4+ and Treg). **B:** *IFNGR1/2* receptor score. **C:** NK and CD8+ T cells in a representative pre-BCG slice. **D:** CXCL9/10/11 score by neighboring T cell infiltration in pre-BCG tumor Visium dots only.

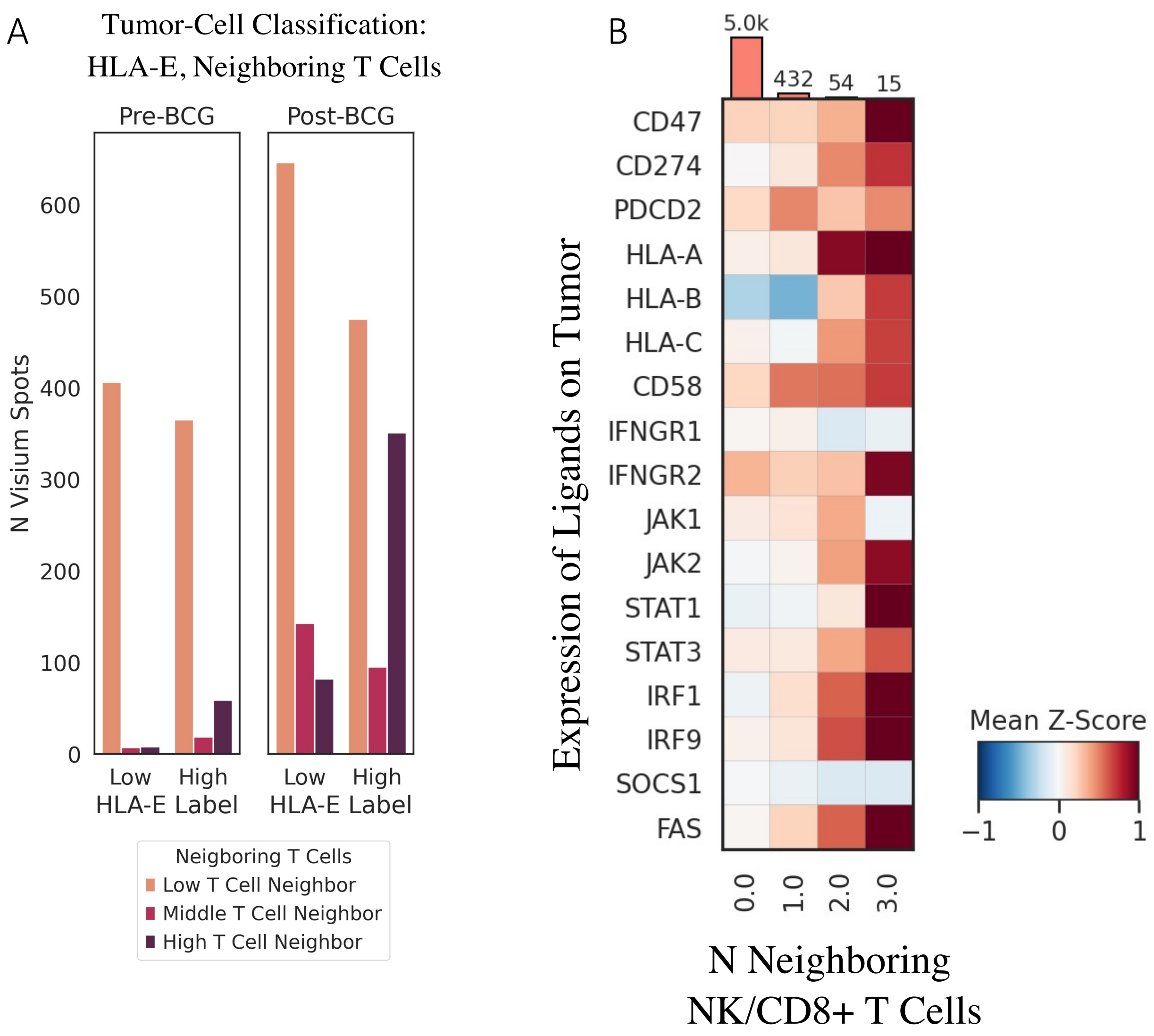


**Supplementary Figure S4:** Spatial neighbors analysis on NK / CD8+ T cells. **A:** tumors classified by HLA-E status (low < 25$^{th}$ percentile; high > 75$^{th}$ percentile) and neighboring CD4+ T cell infiltration pattern. **B:** meta-analysis of tumor Visium spots shown as a function of neighboring NK and CD8+ T cells.

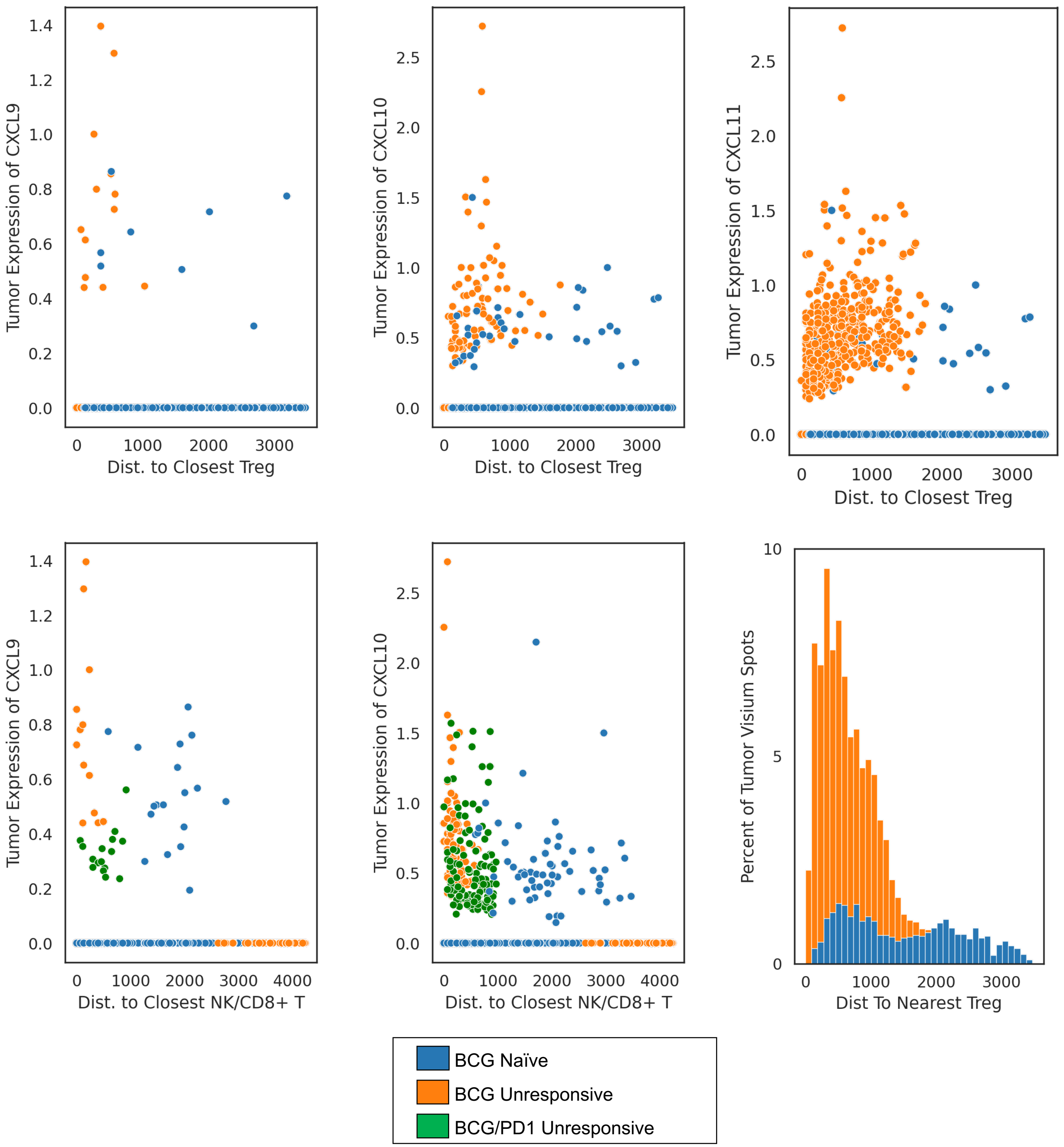


**Supplementary Figure S5:** Spatial neighbors analysis showing the distance between each tumor-identified Visium spot and the closest neighboring cytolytic (NK or CD8+ T cell) or Treg.

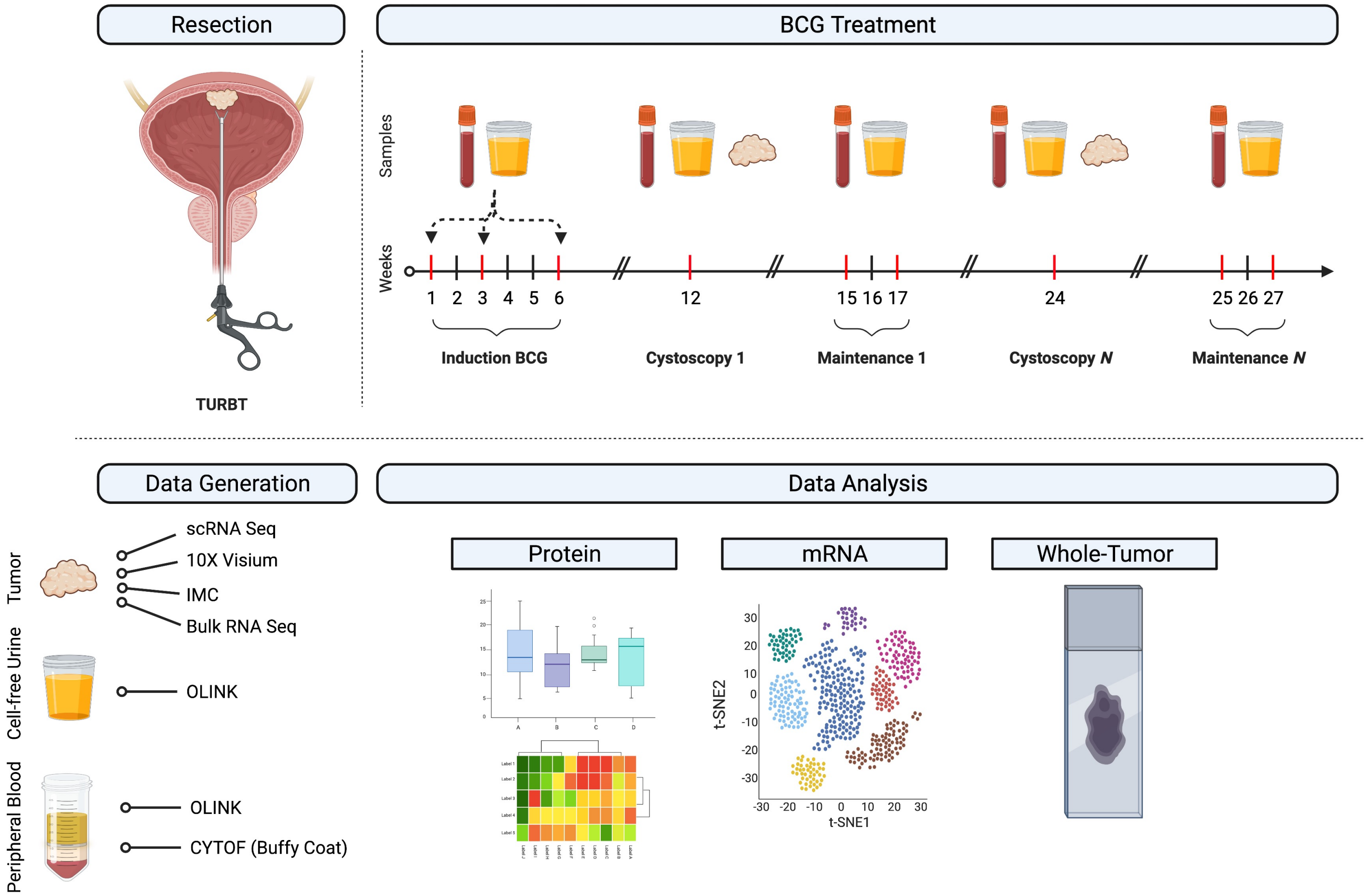


**Supplementary Figure S6:** Overview figure showing BCG treatment and timepoints of sample collection.
